# Functional Resilience of the Neural Visual Recognition System Post-Pediatric Occipitotemporal Resection

**DOI:** 10.1101/2024.05.08.592792

**Authors:** Michael C. Granovetter, Anne Margarette S. Maallo, Shouyu Ling, Sophia Robert, Erez Freud, Christina Patterson, Marlene Behrmann

## Abstract

In the typically developing (TD) brain, neural representations for visual stimulus categories (e.g., faces, objects, and words) emerge in bilateral occipitotemporal cortex (OTC), albeit with weighted asymmetry; in parallel, recognition behavior continues to be refined. A fundamental question is whether two hemispheres are necessary or redundant for the emergence of neural representations and recognition behavior typically distributed across both hemispheres. The rare population of patients undergoing unilateral OTC resection in childhood offers a unique opportunity to evaluate whether neural computations for visual stimulus individuation suffice for recognition with only a single developing OTC. Here, using functional magnetic resonance imaging, we mapped category selectivity (CS) and neural representations for individual stimulus exemplars using repetition suppression (RS) in the non-resected hemisphere of pediatric OTC resection patients (*n* = 9) and control patients with resection outside of OTC (*n* = 12), as well as in both hemispheres of TD controls (*n* = 21). There were no univariate group differences in the magnitude of CS or RS or any multivariate differences (per representational similarity analysis) in neural activation to faces, objects, or words across groups. Notwithstanding their comparable neural profiles, accuracy of OTC resection patients on face and object recognition, but not word recognition, was statistically inferior to that of controls. The comparable neural signature of the OTC resection patients’ preserved hemisphere and the other two groups highlights the resilience of the system following damage to the contralateral homologue. Critically, however, a single OTC does not suffice for normal behavior, and, thereby, implicates the necessity for two hemispheres.

## Introduction

The recognition of visual stimuli is chiefly governed by bilateral neural representations in posterior occipitotemporal cortex (OTC)^1–3^. Disparate but overlapping regions within OTC are implicated in deriving neural representations of images from categories such as faces, objects, and words^4,5^. The development of this brain organization and associated visual competence emerge slowly and dynamically over childhood^6^, likely driven by plastic processes that are more common at younger ages^7,8^ and continue to be refined even to age 30 years^9^.

Functional magnetic resonance imaging (fMRI) studies in typically developing (TD) children attest to the changes in the strength and topography of neural representations for visual categories over time. For example, relative to adults, children show diminished neural activation for faces, larger variability in the spatial location of this activation^10^, and a greater degree of non-specific activation of cortex outside OTC even when just passively viewing face stimuli^11^.

Furthermore, with reading acquisition, competition for cortical territory results in the “recycling” of cortex previously selective, perhaps weakly, for other stimuli to become selective for both words and faces. Initially faces are represented bilaterally, but with the acquisition of literacy, the left and right hemispheres ultimately develop biases for words and faces, respectively^12–15^. Collectively, these studies indicate that the mature pattern of neural representations is a product of cooperative and competitive plastic processes that occur over development, with category-selective neural representations for faces and words−as well as objects^16^−becoming more reliably localizable and precise with experience^6^. This previous work also highlights the fact that, while visual category representations become segregated between the two hemispheres by adulthood, specialization of the two hemispheres in early childhood is not as well-defined^6,12,17^.

The question we address here concerns the malleability of this brain organization during childhood and its sufficiency for visual recognition, and while we examine this in the context of visual cortex, the findings may also have implications for plasticity and/or resilience of neural systems across the brain. Individuals who have undergone surgical removal of OTC in one hemisphere for the management of pediatric drug-resistant epilepsy (DRE) provide a unique opportunity to investigate topographical adaptivity when all category-selective neural responses must coexist in the single preserved OTC. Because plasticity is more viable in childhood than in adulthood, this investigation may permit inferences about topographic malleability and the relationship between the neural profile and behavior^18,19^. Remarkably, fMRI studies in children with a single OTC indicate that the magnitude and location of peak activation is comparable to that observed in age-matched TD controls^20–22^. Moreover, these patients’ multivariate response pattern of category-selective activation across voxels within a region of interest (ROI) do not differ from that of TD age-matched controls, as depicted using category-based representational similarity analysis (RSA)^21,23^. However, our day-to-day visual recognition depends not on differentiating between categories but, rather, on the ability to discriminate among exemplars *within* a category (for instance, recognizing the face of one’s parent versus that of one’s teacher). Here, we examine whether, in a single hemisphere, representations permitting stimulus individuation are also preserved. If the neural signatures for stimulus individuation are unimpaired in pediatric OTC resection patients relative to age-matched controls, such a demonstration would attest to remarkable resilience of cortical organization with a single intact OTC.

The extent to which stimuli are individuated can be gauged by measuring repetition suppression (RS), defined as the reduction of neural activity within a category-selective region of interest (CS ROI) with presentation of a sequence of identical stimuli compared with a sequence of non-identical stimuli^24,25^. Upon repeated presentation of the same stimulus, the observed hemodynamic response is driven lower to a state of habituation or adaptation (fewer neurons are recruited to respond and/or previously active neurons enter a refractory state)^26^. On the other hand, with successive presentation of unique stimuli, the observed overall hemodynamic signal remains relatively high. By measuring RS, we can infer a neural system’s ability to differentiate individual stimuli within a category^27,28^: if a voxel shows RS, this indicates that neurons within that voxel selectively are tuned for the individual token. This exemplar- specificity cannot be inferred by the standard practice of averaging the hemodynamic response signal across all voxels and allows a more fine-grained assessment of the neural system at subvoxel resolution^25^.

The neural processes underlying stimulus individuation appear to emerge over childhood once the category selectivity (CS) profile is established: adolescents, but not children, show reliably localizable CS for faces and scenes in OTC, but neither group shows adult-like RS to individual faces or scenes^29^. That patients with resections that include occipital and/or temporal cortex have seemingly intact CS ROIs^20–22^ does not necessarily imply that the neurons in these regions are tuned for stimulus exemplars. Indeed, while patients with a single OTC perform surprisingly well on recognizing individual faces and words (average accuracy about 85%), their exemplar-specific performance is still statistically inferior to TD controls^30–32^, perhaps due to compromised neural profiles for stimulus individuation.

The current investigation compares the neural profile of both CS and RS in patients with a single OTC (“OTC patients”) with that of age-matched control patients who have undergone surgery outside of OTC (including patients with focal anterior temporal resections; “control patients”) and TD controls. For all participants, CS and RS is measured in response to faces, objects, and words, and participants’ behavioral abilities are characterized as well. If OTC patients do not evince RS in the TD range, this may offer a direct explanation for patients’ previously reported behavioral difficulties^30–32^. If RS is intact but visual behaviors are not, this may suggest that the documented behavioral deficits in visual recognition are less a consequence of impairment of the integrity of stimulus-specific neural representation than they are a consequence of constraining processes to a single hemisphere (and hence, revealing the necessity of a second hemisphere). In addition to quantifying the RS profile of the OTC patients, we also examine whether the RS profiles differ depending on which hemisphere is resected, given the left and right hemispheric specialization for word and face representations that respectively emerge over development^6,29^; this comparison allows us to test whether category and individual visual representations can emerge in either the left or right hemisphere (LH or RH) depending on which is intact and whether one of these hemispheres suffices for behavior that is comparable to controls. Together, the findings will highlight the extent to which the developing, plastic brain subserves changes in or maintenance of the development of neural representations following pediatric cortical resection and the consequences that may bear on behavior.

## Results

This investigation was designed to characterize and compare the neural underpinnings and behavioral performance of face, object, and word recognition in OTC patients, control patients, and TD controls. See Figure 1 and Table 1 for a description of patients. First, for each participant, CS magnitude was quantified for each stimulus category. Second, an fMRI adaptation protocol was deployed to assess neural activation patterns underlying stimulus individuation per category. Note that in the neuroimaging experiments, we are only evaluating patients’ preserved hemisphere. Finally, the participants’ ability to discriminate between exemplars, separately for each stimulus category, was assessed.

**Figure 1:**
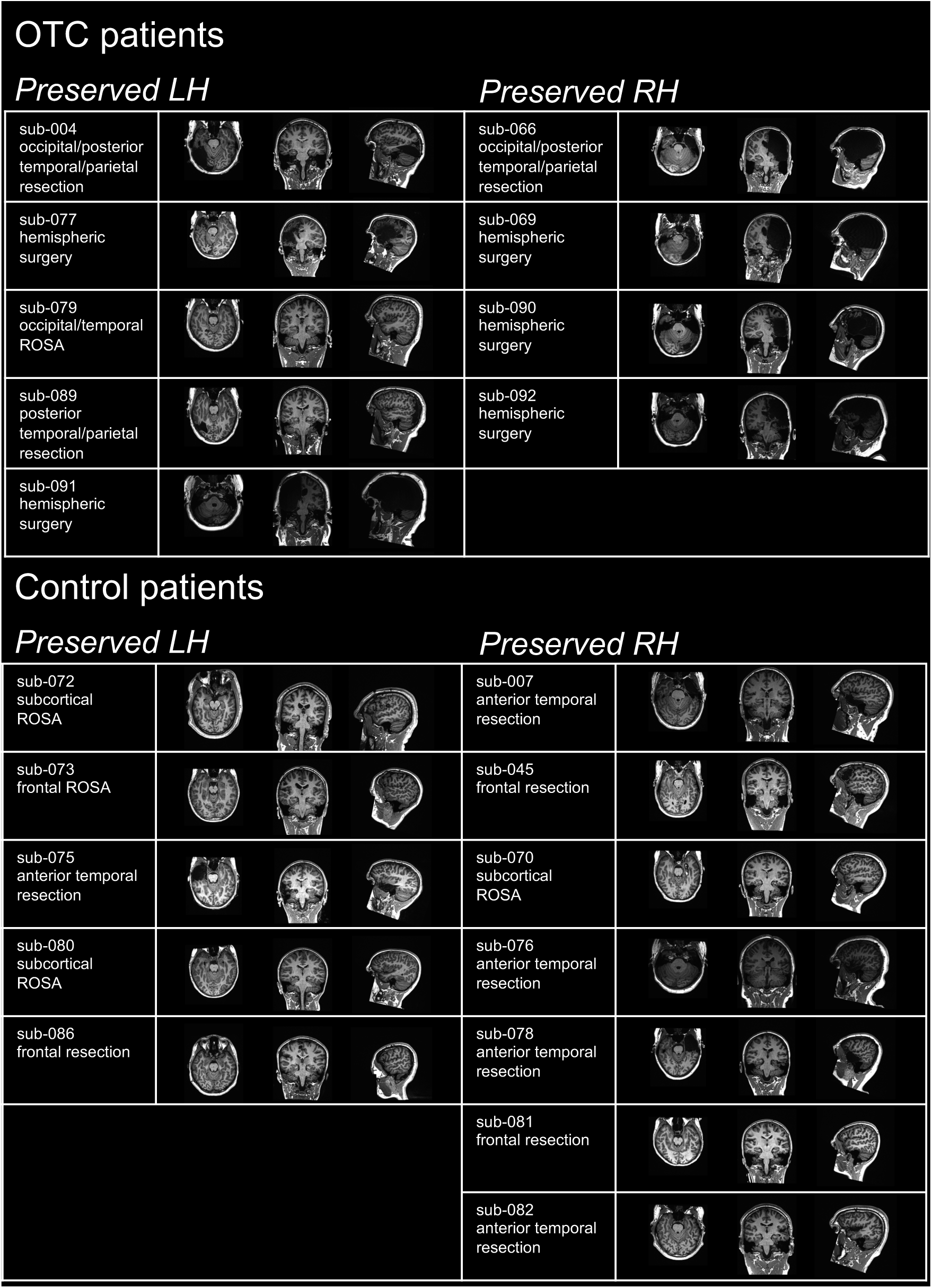
Patient Information. For each participant, sample slices of T1-weighted magnetic resonance imaging are shown in the axial (left), coronal (middle), and sagittal (right) planes at approximately the center of mass of the hemisphere that was the surgical target. Images are shown in radiological convention with the LH on the right and RH on the left. OTC = occipitotemporal cortex. LH = left hemisphere. RH = right hemisphere. ROSA = robot-assisted stereotactic ablation.

**Table 1:**
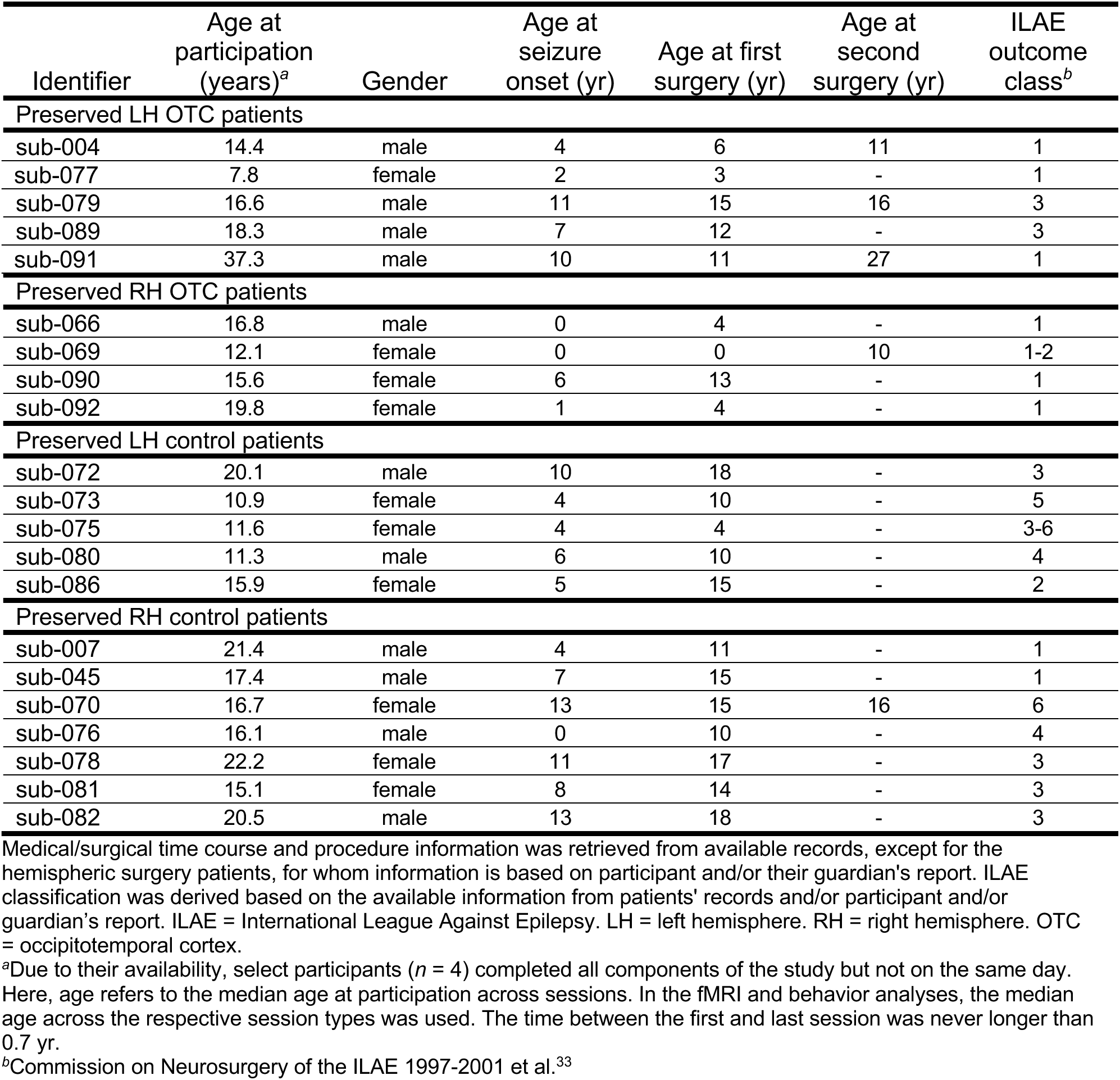
Patient Information.

For each dependent measure, the following statistical procedures are applied: 1) model comparisons of linear mixed effects models (LMEMs) with the likelihood ratio test (LRT) to select the most complex model that minimizes the Akaike information criteria (AIC)^34^, 2) Wald χ^2^-test on the selected model to determine which predictors (e.g., group, hemisphere) are statistically significant^35^, 3) Bayes factor (BF) calculation on each model term to offer evidence either for the null model lacking the factor or for the model containing the factor^36^, and, 4) for statistically significant interaction terms, post hoc contrasts of estimated marginal means (EMM; with the Benjamini-Hochberg correction applied to *p*-values^37^). Additionally, to validate the effect of group on each dependent variable, each model was refit 1000 times with the group label randomly shuffled with replacement, to simulate a random distribution; the *p*-value was taken to be the proportion of instances in which the test statistic in the simulated distribution exceeded the true test statistic^21,38,39^. Last, each patient’s dependent measure was compared to the respective distribution of the TD controls’ values using a procedure for testing differences between individual patients against the control group^40^. Note: mean-centered age was included as a covariate in all models/tests.

### Category Selectivity

Participants viewed blocks of stimuli (faces, objects, words, houses, scrambled objects) while lying in an MRI scanner (Fig. 2a). CS ROIs were defined as clusters of voxels in ventral stream visual cortex (as defined by Glasser et al.^41^) that are significantly more responsive for each primary category of interest in this study (faces, objects, words) vs. a baseline (the other categories). On independent runs of data, CS amplitude was computed as the median responsiveness (β-weight) for a given category vs. baseline of the voxels in that ROI (Fig. 2b).

**Figure 2:**
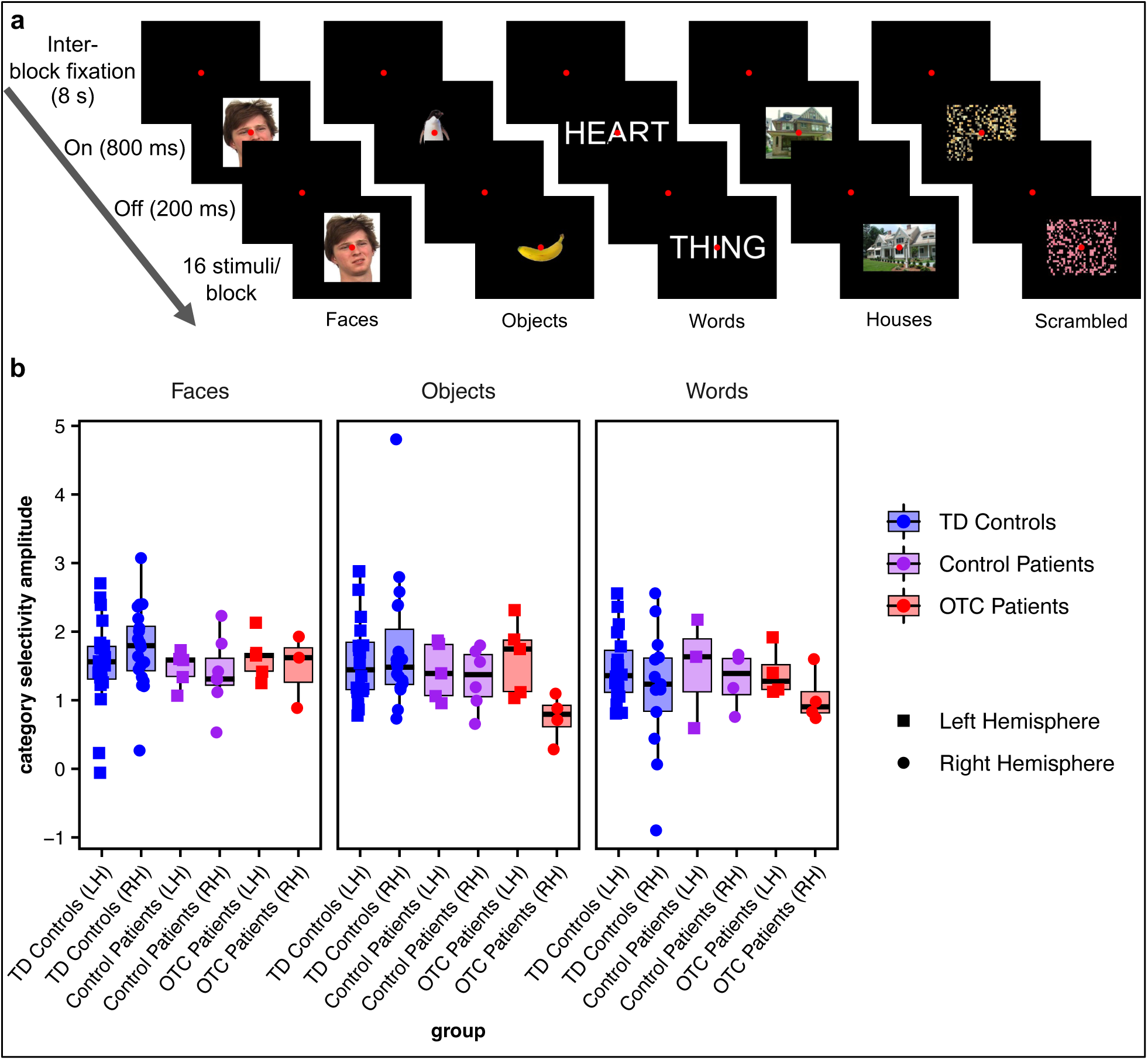
C*a*tegory *selectivity*. (a) Category localizer experiment conducted while participants underwent functional magnetic resonance imaging, adopted from Liu et al.^21^. Face stimuli are from the Face Place dataset^42^. (b) Category selectivity amplitude for each group (boxplots) and individual participant (overlaid, randomly jittered, point plots) by stimulus category (faces, objects, and words). Note that only the patients’ preserved hemisphere data are plotted.

Initially, a LMEM model was fit predicting CS amplitude from the three-way interaction of group (OTC patients vs. control patients vs. TD controls) x hemisphere (LH vs. RH) x stimulus category (faces vs. objects vs. words). No interaction effects survived model selection (see STAR Methods and Table S1 for details). There was only a statistically significant difference in CS amplitude by stimulus category (χ^2^2 = 7.03, *p* = 0.03; BF = 5.41) with faces eliciting a stronger effect than words (for complete post hoc comparison details, see Table S2) but not by group (χ^2^2 = 2.13, *p* = 0.34; BF = 59.02) or by hemisphere (χ^2^1 < 0.01, *p* = 0.97; 12.87; see Table 2 for EMMs by group). Permutation testing confirmed that there was no statistically significant difference in CS amplitude across groups (*p* = 0.37). Additionally, on individual case-control comparisons, no patient (OTC or control) lay outside of the TD control distribution in CS amplitude for any stimulus category in either hemisphere (see Table S3 for statistics). Thus, altogether, patients’ CS amplitude in the contralesional hemisphere is comparable to that of TD controls, independent of whether the LH or RH OTC was resected, replicating prior findings^20,21^.

**Table 2:** Estimates of Dependent Measures.

|  | CS<br>Amplitude | Responsivity:<br>Adaptation<br>Experiment | RS<br>Magnitude | RSA<br>Model Fit | Behavioral<br>Accuracy |
| --- | --- | --- | --- | --- | --- |
| TD<br>Controls | 1.54<br>(1.36 - 1.72) | Faces<br>0.74<br>(0.63 - 0.86) | 0.27<br>(0.22 - 0.33) | 0.76<br>(0.71 - 0.81) | Faces<br>0.90<br>(0.85 - 0.95) |
|  |  | Objects<br>0.54<br>(0.43 - 0.65) |  |  | Objects<br>0.79<br>(0.74 - 0.84) |
|  |  | Words<br>0.82<br>(0.70 - 0.94) |  |  | Words<br>0.96<br>(0.92 - 1.01) |
| Control<br>Patients | 1.35<br>(1.08 - 1.63) | Faces<br>0.74<br>(0.57 - 0.90) | 0.26<br>(0.18 - 0.35) | 0.76<br>(0.68 - 0.84) | Faces<br>0.90<br>(0.84 - 0.95) |
|  |  | Objects<br>0.43<br>(0.25 - 0.60) |  |  | Objects<br>0.71<br>(0.66 - 0.77) |
|  |  | Words<br>1.03<br>(0.84 - 1.23) |  |  | Words<br>0.96<br>(0.90 - 1.02) |
| OTC<br>Patients | 1.31<br>(1.00 - 1.62) | Faces<br>0.73<br>(0.53 - 0.93) | 0.24<br>(0.15 - 0.33) | 0.76<br>(0.67 - 0.85) | Faces<br>0.73<br>(0.67 - 0.80) |
|  |  | Objects<br>0.47<br>(0.28 - 0.67) |  |  | Objects<br>0.55<br>(0.47 - 0.62) |
|  |  | Words<br>0.75<br>(0.56 - 0.95) |  |  | Words<br>0.93<br>(0.86 - 0.99) |

Estimated marginal means (95% confidence interval). Estimates are computed per stimulus category for responsivity in the adaptation experiment and for behavioral accuracy because there was a significant interaction effect of group x stimulus category on both dependent measures. CS = category selectivity. RS = repetition suppression. RSA = representational similarity analysis.

### Repetition Suppression

Participants viewed blocks of stimuli (faces, objects, words) in which the same stimulus was displayed successively (“same” condition), two different stimuli were shown successively alternating (“alternate” condition), or each stimulus displayed in succession was unique (“different” condition; Fig. 3a). Per participant, a generalized linear model (GLM) was fit to the fMRI data, and the median β-weight for a block type (e.g., same faces, alternating objects, etc.) across voxels within the “preferred” CS ROI was considered the “responsivity” to that block type (Fig. 3b).

**Figure 3:**
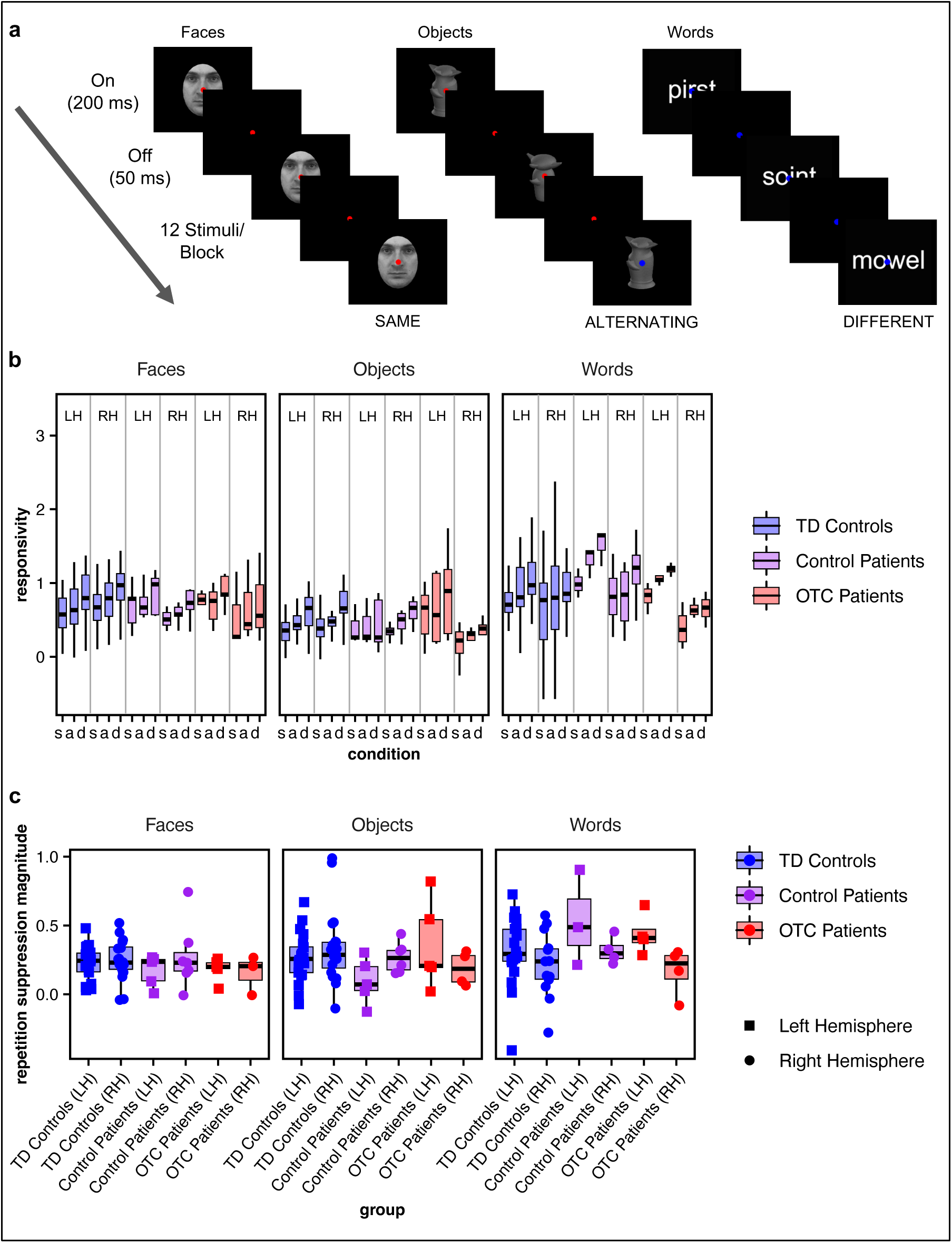
R*e*petition *suppression*. (a) Adaptation experiment conducted while participants underwent functional magnetic resonance imaging. Face stimuli are from the Karolinska Directed Emotional Faces stimulus set (stimulus ID AM03NES in the shown example)^43^. (b) Responsivity for each group per condition (same, alternating, different) by stimulus category (faces, objects, and words; boxplots). (c) RS magnitude (e.g., the contrast of same versus different blocks) for each group (boxplots) and individual participant (overlaid, randomly jittered point plots) by stimulus category (faces, objects, and words). Note that only patients’ preserved hemisphere data are plotted.

Initially, a LMEM was fit to test the four-way interaction of block condition (same, alternating, different) x group x hemisphere x stimulus category on the responsivity within the corresponding CS ROI. The four-way interaction did not survive the model selection (see Table S4 for details), and there were also no significant three-way interactions (group x condition x stimulus category: χ^2^8 = 2.57, *p* = 0.96, BF = 1.71×10^10^; group x hemisphere x stimulus category: χ^2^4 = 5.04, *p* = 0.28, BF = 2.03×10^4^; group x condition x hemisphere: χ^2^4 = 0.87, *p* = 0.93, BF = 1.61×10^5^; condition x hemisphere x stimulus category: χ^2^4 = 1.29, *p* = 0.86, BF = 1.30×10^5^). Additionally, although there was a significant two-way interaction of group x stimulus category (χ^2^4 = 10.70, *p* = 0.03), there is strong evidence favoring the null model without the interaction term (BF = 1.37×10^3^). Post hoc comparisons showed that there were no differences between groups for any stimulus category, which was confirmed with permutation testing (see Table 2 for EMMs by group x stimulus category and Table S5 for complete post hoc comparison details). There was also a significant two-way interaction effect of hemisphere x stimulus category (χ^2^4 = 12.44, *p* < 0.01, BF = 1.20; see Table S6 for post hoc comparisons) but no other significant two-way interaction effects (group x condition: χ^2^4 = 0.15, *p* :: 1, BF = 2.29×10^5^; group x hemisphere: χ^2^2 =4.76, *p* = 0.09, BF = 51.06; condition x hemisphere: χ^2^2 = 0.03, *p* = 0.99, BF = 4.87×10^2^; condition x stimulus category: χ^2^4 = 1.22, *p* = 0.88, BF = 1.37×10^5^). Additionally, there was a significant effect of condition (χ^2^2 = 58.82, *p* < 0.001, BF = 2.60×10^-9^). Post hoc testing demonstrated that the responsivity to the different condition was significantly greater than that for each the alternating condition (*z* = 3.43, *p* < 0.001) and same condition (*z* = 6.04, *p* < 0.001), and the responsivity to the alternating condition was significantly greater than that for the same condition (*z* = 2.61, *p* = < 0.001), as would be expected with RS^24,25,28^, thereby validating the study paradigm. Finally, on individual case-control comparisons, in all but one case (a single right-sided resection OTC patient’s responsivity to different objects in the LH vs. TD controls’ LH), there were no significant differences between any patient and the TD controls (see Table S7 for statistics).

Next, to examine RS specifically, we computed the difference per voxel in the responsivity for a different block and same block within the predefined CS ROI associated with the stimulus class being presented in the given blocks. We defined the RS magnitude as the median value across voxels (Fig. 3c). Initially a LMEM was fit with the three-way interaction of group x hemisphere x stimulus category. Only the three-way interaction did not survive model selection (see Table S8 for details). There was a significant two-way interaction effect of hemisphere x stimulus category (χ^2^2 = 9.04, *p* = 0.01, BF = 2.05). Post hoc comparisons revealed that RS to words in the LH was significantly greater than RS to faces in the LH (*z* = 3.84, *p* < 0.01; as expected on accounts of weighted asymmetry for words in LH^6^), RS to objects in the LH (*z* = 3.15, *p* = 0.01), and RS to words in the RH (*z* = 2.51, *p* = 0.04), and there were no other significant pairwise differences (see Table S9 for complete post hoc comparisons). The two-way interaction of group x hemisphere (χ^2^2 = 2.51, *p* = 0.28, BF = 48.65) and group x stimulus category (χ^2^4 = 8.45, *p* = 0.08, BF = 4.48×10^2^) on RS magnitude were not statistically significant. As with CS amplitude, and concordant with our analyses on responsivity to the adaptation experiment above, there was no significant difference in RS magnitude by group (χ^2^2 = 0.34, *p* = 0.84, BF = 3.41×10^6^; see Table 2 for EMMs), as further confirmed by permutation testing (*p* = 0.86). Moreover, on individual case-control comparisons, in all but one case (a LH resection control patient with lower RS magnitude to faces in the RH vs. TD controls’ LH), there were no significant differences between any patient from the OTC or control patient group and TD controls (see Table S3 for statistics).

Thus altogether, as with CS, there were no group differences in RS between OTC patients and control patients or TD controls.

### Representational Similarity Analysis

To scrutinize further the extent to which the neural representations of the OTC patients match those of the other two groups, RSA was conducted. RSA involves computing a second-order correlation (e.g., Spearman’s correlation) between activation-pattern representational dissimilarity matrices (RDMs) and model RDMs. Model RDMs represent the predicted similarity between experimental conditions based on a priori hypotheses, and activation-pattern RDMs are computed from the neural profile using dissimilarity metrics (e.g., Pearson’s correlation). Fig. 4a displays the average activation RDMs on the fMRI adaptation data for each participant group (note patients’ data are collapsed across hemispheres here due to small sample sizes and, as above, we know that there are no group by hemisphere differences). Specifically, each participant’s activation RDM was transformed to Fisher’s *z*-score, averaged within the group, and then transformed back to Pearson’s correlation values. To determine the similarity of the RDMs across groups, the rank correlation (Kendall’s *τ*) of each pair of RDMs was computed.

**Figure 4:**
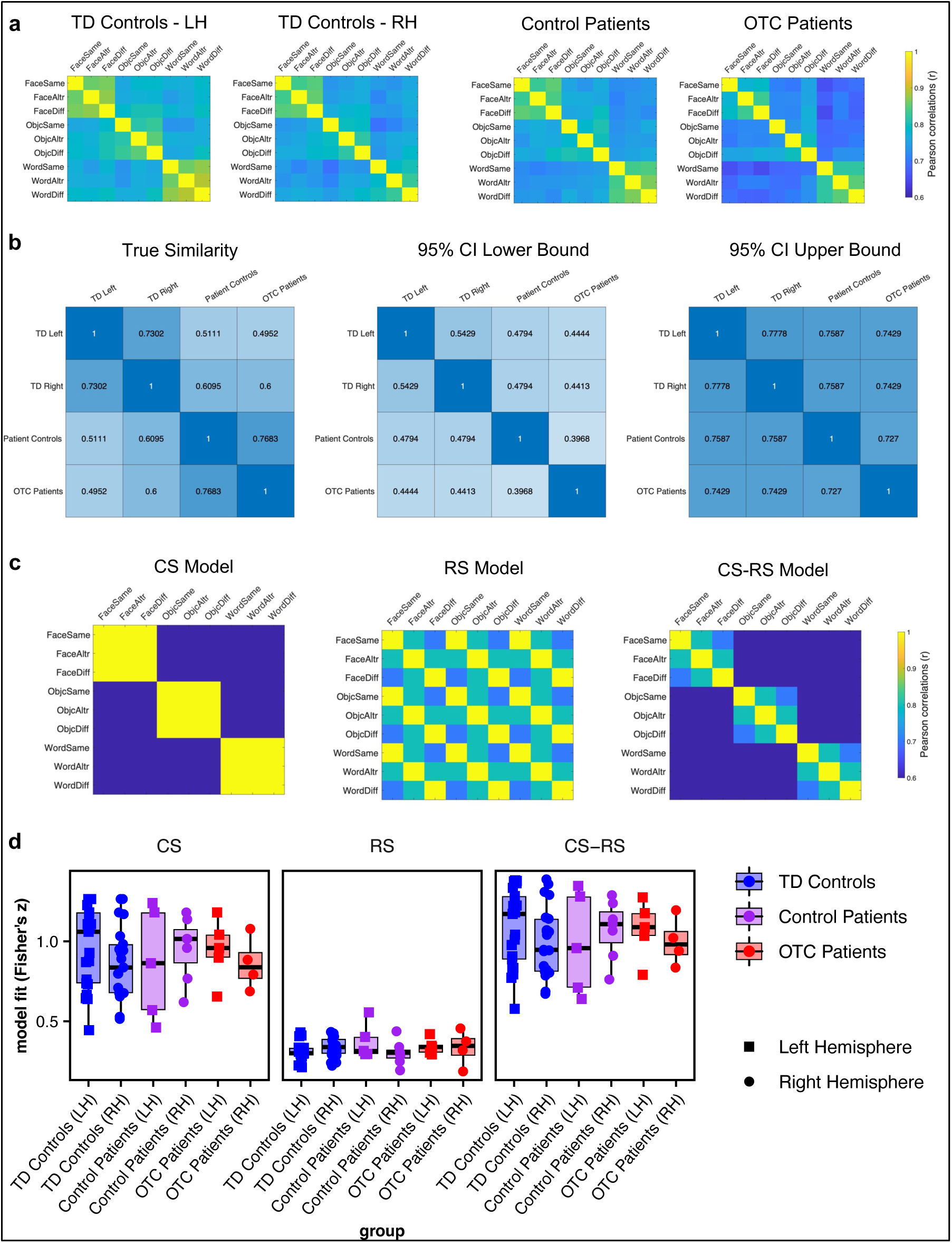
R*e*presentational *similarity analysis*. (a) RDMs per group (note patient groups are collapsed across hemisphere here due to small sample sizes per group). (b) Permutation testing was used to simulate a null distribution assuming groups are like one another; the true similarity matrix on the far left lies within the lower bound (middle) and upper bound (far right) of the simulated distribution. (c) Theoretical models for CS, RS, and a combination of CS and RS. (d) Model fit for each group (boxplots) and individual participant (overlaid, randomly jittered point plots) by model type (CS, RS, CS-RS). Note that data only from the patients’ preserved hemisphere are plotted.

Then, over 5000 iterations, group labels were shuffled, group average RDMs were computed per group, and pairwise correlations were calculated. All participant groups exhibited similar patterns of activation (see Fig. 4b).

We then correlated each participant’s RDM with three theoretical models, using, in an assumption-free approach, all ventral stream visual cortex voxels, which is narrowly defined as just posterior regions by the Glasser atlas, on the fMRI adaptation data (Fig. 4c). First, a CS model was designed where ventral cortex exhibits CS without RS, with activation patterns across conditions of the same category (same, alternating, or different) showing high similarity, while patterns of different categories showing low similarity. Second, a RS model was constructed, in which voxels show RS but do not distinguish between categories. We theorized that the similarity in a same vs. different comparison would decrease compared to a same vs. same comparison, illustrating the classic RS effect; the similarity in a same vs. alternating comparison was hypothesized to be intermediate. Additionally, we expected that the similarity in an alternating vs. different comparison would fall in between a same vs. same comparison and a same vs. different comparison. Last, a CS-RS model combined both RS and CS, such that RS should be present within a category but not across categories.

A LMEM was initially run predicting model fit (Fischer’s *z*) from the three-way interaction of group (TD controls, control patients, OTC patients) x hemisphere (LH, RH) x model type (CS, RS, CS-RS; Fig. 4d). No interaction effects survived model selection (see Table S10). There was a significant difference in model fit across the three models (χ^2^2 = 807.23, *p* < 0.001, BF = 7.57×10^60^); post hoc contrasts indicated that the CS model fit significantly better than the RS model (*z* = 21.91, *p* < 0.001). The CS-RS model, however, was the best fit for the data, significantly outperforming the CS model (*z* = 4.71, *p* < 0.001) and the RS model (*z* = 26.62, *p* < 0.001). Importantly, there was no effect of group (χ^2^2 = 0.03, *p* = 0.99, BF = 1.86×10^2^), nor of hemisphere (χ^2^1 = 3.71, *p* = 0.05, BF = 2.22) on the model fit. Permutation testing confirmed that there was no difference in model fit across groups (*p* :: 1). Moreover, individual case-control comparisons demonstrated that no patient’s model fit fell outside the distribution of model fits for each the LH and RH of TD controls, except for the RS model for one right-resection control patient compared to the distribution of model fits for TD controls’ LH (see Table S11 for statistics).

Overall, the multivariate approach to examining neural activation pattern in OTC patients, control patients, and TD controls revealed no group differences.

### Behavior

In addition to undergoing neuroimaging, participants completed tasks requiring discrimination between each of faces (Fig. 5a), objects (Fig. 5b), and words (Fig. 5c). Data are plotted in Fig. 5d.

**Figure 5:**
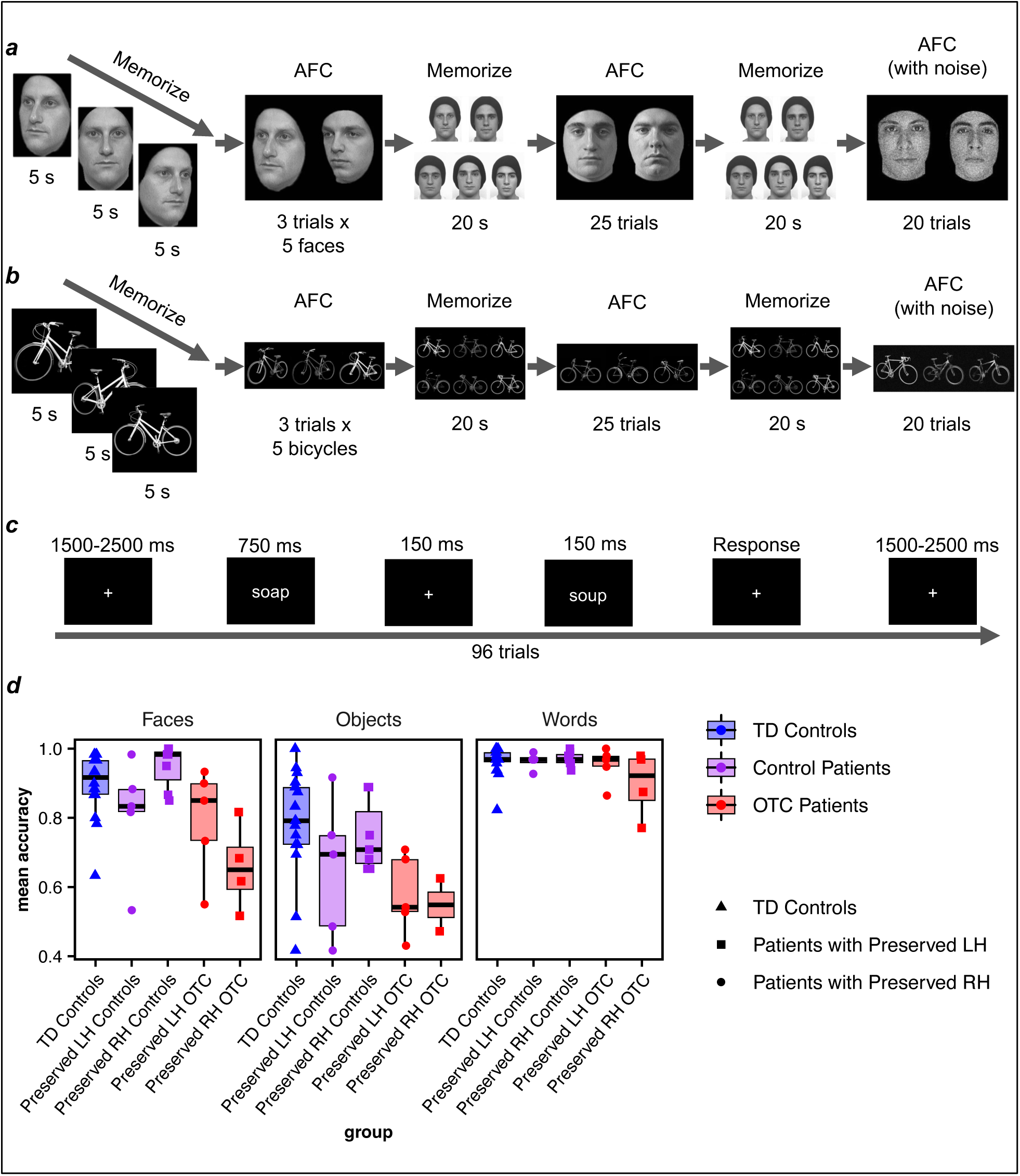
B*e*havior. Designs of the (a) Cambridge Face Memory Test^44,45^ (figure adapted from Croydon et al.)^44^, (b) Cambridge Bicycle Memory Test^46,47^ (figure adapted from Dalrymple et al.)^47^, and (c) word recognition task^30,48^ (figure adapted from Dundas et al.)^48^. (d) Mean accuracy for each group (boxplots) and individual participant (overlaid, randomly jittered point plots) by task (recognition of faces, objects, words).

A LMEM (see Table S12 for model selection details) was fit predicting task accuracy from the interaction effect of group (TD controls, control patients, OTC patients) x stimulus category (faces, objects, words; Fig. 5b); this interaction effect was significant (χ^2^4 = 17.62, *p* < 0.01, BF = 6.25). Group comparisons demonstrated that OTC patients’ face recognition accuracy was significantly lower than that of both TD controls (*z* = 4.06, *p* < 0.001) and control patients (*z* = 3.67, *p* < 0.001). Control patients and TD controls did not differ in performance (*z* = 0.10, *p* = 0.92), suggesting that the group deficit observed among OTC patients was specific to resection inclusive of OTC. On object recognition, OTC patients’ accuracy was also significantly lower than that of both TD controls (*z* = 5.37, *p* < 0.001) and control patients (*z* = 3.45, *p* < 0.01). Again, control patients and TD controls did not differ in performance (*z* = 2.02, *p* = 0.08), suggesting that the deficit was specific to OTC patients. There were no group differences on word recognition (see Table 2 for EMMs and Table S13 for complete post hoc comparison statistics). Permutation testing confirmed group comparisons (Table S13).

As TD controls have two intact hemispheres, hemisphere could not be modelled as a predictor for accuracy; therefore, an analysis was performed only on the OTC and control patients’ data. Initially, a LMEM predicting accuracy from the three-way interaction of group (control patients, OTC patients) x hemisphere (LH, RH) x stimulus category (faces, objects, words) was fit on only the patients’ data. The three-way interaction did not survive model selection (see Table S14 for details), but there were significant interaction effects of both group x hemisphere (χ^2^1 = 4.32, *p* = 0.04, BF = 1.03) and group x stimulus category (χ^2^2 = 7.42, *p* = 0.02, BF = 2.91) on accuracy; there was no interaction effect of hemisphere x stimulus category (χ^2^2 = 0.13, *p* = 0.94, BF = 56.30). As above, relative to control patients, OTC patients had significantly lower accuracy on face recognition (*z* = 3.42, *p* < 0.01) and object recognition (*z* = 3.22, *p* < 0.01) but not word recognition (*z* = 0.57, *p* = 0.57). Decomposing the group x hemisphere interaction revealed that left OTC resection patients had significantly lower accuracy than left resection control patients (*z* = 3.83, *p* < 0.01), but otherwise there were no further pairwise differences (see Table S15 for complete post hoc statistics).

Finally, on individual comparisons of each OTC and control patient to the TD control distribution, three out of nine OTC (two preserved LH, one preserved RH) and only one out of 12 control patients (preserved RH) showed significant deficits on face recognition. On word recognition, one out of nine OTC patients (preserved LH) and no control patients exhibited a deficit. There were no observed deficits on object recognition (see Table S3 for statistics). Of note, all OTC patients performed all tasks at above-chance accuracy.

Altogether, the findings imply that OTC patients have significantly lower accuracy on both face and object−but not word−recognition, and that this deficit appears specific to surgery that encompasses OTC.

## Discussion

Here, we examine the nature of neural representations underlying visual category recognition and stimulus individuation in participants who, following childhood resection of OTC for the treatment of DRE, develop with only a single OTC. The neuroimaging experiments revealed several findings, supported by both univariate and multivariate analyses. First, consistent with prior work^20,21^, pediatric OTC patients showed CS amplitudes comparable to that of TD controls and to control patients (with surgeries outside of OTC). This suggests that despite developing with just a single OTC, patients’ contralesional cortex shows differentiable neural representations for different visual stimulus categories, comparable to those representations observed in the developing LH and RH in TD individuals. Second, there were no between-group differences in the magnitude of RS to individual stimulus exemplars measured *within* the contralesional CS ROI. RS is often used to characterize the function of perceptual systems in the brain^26^, both for assessing the extent of representational overlap between exemplars and for inferring the neural computations giving rise to cognitive behaviors^24^. Suppression of the hemodynamic response on fMRI is posited to reflect neural subpopulations’ individual sensitivities to the repetition of exemplars, either through neural “fatigue” from the selective neurons’ delayed refractory periods or neural “tuning” such that neurons that do not represent an exemplar’s features are suppressed to conserve metabolic resources^26^. Our finding of comparable RS across groups thus suggests that the neural processes subserving individuation between exemplars of the same category are intact in patients’ contralesional OTC.

At the same time, despite the apparent integrity of the neural profile, the OTC patients’ visual recognition ability is statistically inferior to that of controls for face and object recognition, as demonstrated previously^30–32^. OTC patients’ behavioral profiles appear to reflect that visual recognition may stringently require the maintenance of representations across two hemispheres to render typical recognition, or in other words, two hemispheres are better than one.

The competitive interplay underlying the establishment of visual stimulus category representations across the two hemispheres over development has long led to the question of whether both hemispheres are necessary for neural representations to mature sufficiently and optimally. Neuropsychology studies in adults reveal that, at later ages, accommodation of new functions is limited, as even a focal unilateral lesion to left or right OTC in adulthood can result in pronounced recognition deficits for words and faces, respectively^49–51^, that are only marginally improved with intervention^52,53^. At the same time, given that competition for representing visual stimulus categories is dynamic and ongoing throughout development^6,12^, it is reasonable to conjecture that representations for the full array of visual stimulus categories may come to co- exist in a single, developing OTC. Indeed, our prior work demonstrated that patients with only one hemisphere could nonetheless perform visual recognition tasks at above-chance accuracy (on average ∼85%) but still poorer relative to healthy controls. The remaining question was could we observe neural signatures suggesting that representations for stimulus individuation can co-emerge in a single OTC? Or does constraining representations to a single hemisphere impair the neural computations for stimulus individuation? And what are the consequences for behavior?

Our results demonstrate that following pediatric OTC resection, neural processes of typical development do not appear to have been interrupted, as OTC patients evince CS and RS profiles that are within the range of that of TD controls. Notably, as there is no interaction of group by hemisphere on either CS or RS, this is the case independent of which hemisphere is resected. These findings highlight the non-resected OTC’s resilience to damage to its contralateral homologue and stability of the typical neurodevelopmental trajectory in that hemisphere. At the same time, that pediatric OTC surgery patients show intact CS ROIs with intact RS, but impaired behavior, suggests that their behavioral impairment is likely not caused by an atypicality in the neural computations underlying stimulus individuation per se (at least at the resolution that RS paradigms can afford). In other words, one *functional* hemisphere may be sufficient−as all OTC patients performed all tasks at above-chance accuracy−but nonetheless suboptimal for visual recognition. The lack of support from the resected hemisphere and/or constraining of function to a single hemisphere may be implicated in the behavioral deficit.

Importantly, we do not observe a difference in CS, RS, or behavioral performance between OTC patients and controls that is a function of the hemisphere resected. This is consistent with findings from other work suggesting that hemispheric specialization is not well-refined early in development. For example, insult to either the LH or RH in early childhood can result in a face recognition deficit^54^, but in adulthood, prosopagnosia is more common and more severe after RH than LH damage^55^. Similarly, there is more bilateral activation in language regions in children compared with adults whose neural profile is strongly weighted to the LH^56^.

One problem with our account arguing for the necessity of two hemispheres for typical behavior is that it does not explain the specific finding in which face and object recognition, but not word recognition, are affected in the OTC cases. An explanation for this finding is that in TD individuals, only one (the left) hemisphere is ultimately more specialized for word recognition, whereas representations for faces and objects are more distributed across the two. Thus, since word recognition relies on only one hemisphere in the TD population^6^, it may be unsurprising that patients with only one intact hemisphere here show no deficits in word recognition.

Moreover, while the OTC patients collectively do not have a specific deficit in word recognition, there is nonetheless a group by hemisphere interaction on task accuracy that is independent of stimulus type. This interaction is accounted for by the fact that left OTC patients perform poorer on visual recognition tasks overall relative to left control patients (a difference not observed between right OTC patients and right control patients). In typical development, word representations largely emerge in the LH, so the finding here suggests that despite the typical neural profile in the RH, the potential load placed on the RH to adopt function not typically ascribed to it may explain the overall deficit in visual recognition in our LH OTC resection cases.

The brain may be resilient to injury at younger ages due to the up-regulation of plasticity. While the current study demonstrates the remarkable stability of one hemisphere forced to develop in isolation, future work is necessary to demonstrate the potential plastic mechanisms subserving this stability. Prior work has demonstrated, for instance, that there are global differences in functional connectivity (FC) in the contralesional hemisphere of this patient population, indicative of plasticity from competition for functional representations^22,57,58^. The mechanisms by which brain-wide FC differences may be implicated in the formation of stimulus exemplar representations remains to be explored. Furthermore, not all neural systems may demonstrate the resilience that we observe here; for example, we know that individuals with resections of parietal cortex show much less plasticity of function to the homologous region of the other hemisphere, unlike in the OTC resections in the current cases^59^.

We should note too that while this study characterizes visual recognition of the largest sample of pediatric OTC patients to date, both in terms of behavioral and neural data, the group of patients was nonetheless heterogenous. This heterogeneity is advantageous in that it suggests that the findings are likely to extend to a diversity of patient cases, and individual case-control comparisons revealed few obvious patterns to suggest that select patient ages or surgery types result in unique behavioral or RS profiles different from the group. At the same time, further study of larger samples of patients would allow one to determine possible patient-control differences dependent on relevant clinical variables including age at disease onset/surgery or medical etiology. Understanding the relationship of age at disease onset/surgery and behavioral/CS/RS profiles will be especially important given the slow maturation of visual category representations that occur over development^10,12,29^. It is possible perhaps that while pediatric OTC surgery patients overall do not show differences from controls, that within this group of patients, those with earlier neural damage may show evidence for greater plasticity than those with later surgeries. Indeed, earlier surgery has been posited to “correct” the developmental trajectory of a child with DRE, as evidenced by improved cognitive outcomes following resection^60^. It will also be critical for future studies to characterize the nature of neural representations for visual recognition pre- and post-surgery to ascertain whether there are differences in plasticity/resilience as a function of disease etiology vs. surgical intervention.

Altogether, this study demonstrates that patients with pediatric OTC surgery show, relative to control patients and TD controls, deficits in visual recognition behaviors but, remarkably, no differences in CS or RS to visual stimulus exemplars in the contralesional, preserved hemisphere. The findings indicate that while the processes of establishing representations for visual stimuli associated with development may not depend on the presence of two hemispheres and are resilient in the context of early injury to one hemisphere, because these processes are constrained to a single hemisphere, behavior may be adversely impacted. That is, despite the stability of the neural profile of the preserved hemisphere to support visual recognition, the absence of the other suggests both are likely necessary for optimal behavior.

## Methods

### Data and Code Availability

Raw behavior and neuroimaging data, as well as all experiment and original code, will be deposited at KiltHub (Carnegie Mellon University’s repository, hosted on figshare) and be made publicly available upon publication (direct object identifier: 10.1184/R1/25727349). Any additional information required to reanalyze the data reported in this paper is available upon request.

### Study Participant Details

Patients (*n* = 21) with childhood cortical resections or ablations were recruited from the Pediatric Epilepsy Surgery Program at University of Pittsburgh Medical Center Children’s Hospital of Pittsburgh or with assistance from the Pediatric Epilepsy Surgery Alliance (formerly the Brain Recovery Project). Of these patients, nine underwent surgeries encompassing OTC (including six with near to total resection of an entire hemisphere but always encompassing OTC). The remaining 12 patients (referred to throughout as “control patients”) underwent surgeries where OTC was preserved bilaterally (including four with anterior temporal resections). All patients had surgery at age 18 yr or younger. Additionally, native-English-speaking, right-handed individuals who had not undergone prior neurosurgery (referred throughout as typically developing or “TD controls”) were recruited from the local Pittsburgh community. See Table S1 for summary statistics of participants’ demographics, Table 1 for individual patient details, and Fig. 1 for an illustration of the OTC patients’ lesions.

To verify the matching of patients and controls specifically with respect to age at time of testing and gender, multinomial regression models were fit predicting group (OTC patients vs. control patients vs. TD controls), with age and gender in separate models. A permutation test was performed in which predictor values were randomly sampled with replacement, and the model was fit on the permuted data, 1000 times. *p* was calculated as the probability that the absolute value of *z* for the predictor (for distinguishing OTC patients from each TD controls and control patients) was greater than or equal to the absolute value of *z* observed. Age was not a significant predictor for distinguishing OTC patients from either TD controls (*z*4 = 0.18, *p* = 0.88) or control patients (*z*4 = 0.18, *p* = 0.73). Gender also was not a significant predictor for distinguishing OTC patients from either TD controls (*z*4 = 0.64, *p* = 0.51) or control patients (*z*4 = 0.25, *p* = 0.82).

Adult participants or child participants’ guardians gave informed consent, and child participants assented to participate. This study was reviewed and approved by the Institutional Review Boards of Carnegie Mellon University and the University of Pittsburgh.

### Method Details

#### Neuroimaging Experiments

All but four participants’ data were acquired with a Siemens Prisma 3T magnetic resonance imaging (MRI) scanner with a 64-channel head coil. Data from the remaining four participants (all TD controls) were collected with a Siemens Verio 3T MRI with a 32-channel head coil. High-resolution T1-weighted whole-brain anatomical data were acquired with a magnetization rapid gradient-echo sequence: 176 slices, field of view (FOV) = 256 x 256 millimeters, repetition time (TR) = 2300 ms, echo time (TE) = 1.97 ms, and scan time = 5 minutes 21 s.

There were two functional scans, one a set of localizers for the selection of CS ROIs^20–22^ and the other, an adaptation paradigm. On each run of the localizer, participants viewed 15 blocks of stimuli (each separated by approximately 8-s inter-block periods with no stimulus presentation, including one prior to the start of the first block) displayed with the Psychophysics Toolbox (version 3^61^) in MATLAB. Each block consisted of 16 stimuli (each shown for approximately 800 ms with a 200-ms interstimulus interval) from one of five categories: faces, objects, words, houses, and scrambled objects (Fig. 2a). Of the 16 stimuli per block, 15 were unique, with 1 stimulus repeated randomly per sequence. A fixation cross was always displayed at the screen’s center, and participants were instructed to maintain fixation and press a button on detection of a stimulus repeat. Each category block (totaling approximately 16 s) was shown three times per run, in a pre-determined random sequence used for all participants, and all participants completed three runs. Across each of the estimated 6 min 8 s runs, echoplanar imaging (EPI) data were acquired (69 slices, FOV = 212 x 212 mm, TR = 2000 ms, TE = 30 ms, multiband acceleration factor = 3).

The fMRI adaptation paradigm^27^ consisted of 6 runs, each comprising 36 “mini-blocks” of approximately 3 s each. In each mini-block, 12 stimuli were presented centrally, each for a 250 ms, with 50-ms intervals between stimuli. For each mini-block, three baseline fixation periods without presentation of stimuli—of durations 6, 7.5, and 9 s each—were randomly interspersed. This mini-block design was intended to increase statistical power while limiting the duration that children would be in the scanner. Valid general linear models (GLMs) can be successfully fit to EPI data with this format, including stimulus category mini-blocks that are likewise not all separated by baseline fixation periods^62^.

Blocks featured one of three stimulus classes: unfamiliar faces (see Table S16 for list of stimulus IDs from the Karolinska Directed Emotional Faces stimulus set^43^), Greebles (a novel object stimulus class^63^), or pseudowords (Fig. 3a), all of which were chosen to measure RS without the confound of prior familiarity with specific stimulus exemplars. Furthermore, there were three block conditions: “same” for which the same exemplar was shown 12 times; “alternating” for which two individual stimuli were shown in an alternating pattern, six times each; and “different” for which 12 unique stimuli were shown in succession. Across the experiment, 18 unique exemplars were presented for each stimulus category. The same two and 12 exemplars were used for the alternating and different conditions, respectively. On the other hand, four unique exemplars were used for each same condition within a run, but the same four exemplars were used for the same condition across runs (presented in a randomized order per run). Blocks were pseudorandomized such that each of the nine block types (three stimulus categories x three conditions) were presented once before cycling through the block types again. Moreover, two blocks of the same stimulus category did not immediately follow one another. All participants viewed blocks in an identical sequence.

Participants were instructed to fixate on a dot in the center and to indicate with a button press when the dot changed color (from blue to red or red to blue) to encourage fixation. On each run, the dot changed color only once per block type, at a random time during the block.

The color change occurred in the same sequence of block types across runs, though block types were randomized in their sequence, limiting the predictability of the dot color change. For three out of the four repetitions of the nine block types within a run, the dot changed color only twice, and on the other, three times, totaling nine color changes across the run. The color change timing was identical across participants.

All participants completed six runs. Across each of the estimated 3 min 36 s runs, EPI data were acquired (68 slices, FOV = 192 x 192 mm, TR = 1500 ms, TE = 30 ms, multiband acceleration factor = 3).

#### Behavioral Experiments

Participants were also assessed on their ability to recognize faces, objects, and words. The Cambridge Face Memory Test adapted for children^44^ was used (Fig. 5a). In the first block, participants viewed a grayscale male face sans hair from three different angles, each for 5 s. Then, in an alternative forced choice task, they were presented with three face pairs (each pair shown at one of three angles) successively and identified which of the two faces they had just seen. This process was repeated for five faces, totaling 15 trials. In the second block, participants studied the previously presented five faces for 20 s, after which they were presented with face pairs (one “old” and one “distractor”) and identified which of the two faces was “old” (25 trials). In the final block, after studying the five faces again for 20 additional seconds, participants identified which of the two faces was “old,” but in the final 20 trials, faces were presented with Gaussian noise to increase the difficulty of the task. All participants viewed the same stimuli presented in the same sequential order. Accuracy was computed as the mean accuracy across all trials from the three blocks (60 total).

Participants also completed the Cambridge Bicycle Memory Test with grayscale bicycles instead of faces^46^ (Fig. 5b). The experiment structure was identical to that of the Cambridge Face Memory Test, except that participants were initially presented with and subsequently studied a total of six, not five, exemplars; and participants identified the exemplar they had previously seen among three, not two, stimuli.

Participants additionally completed a word matching task, previously used in this patient population^30^. Pairs of 58 four-letter gray words, chosen to be age-appropriate for young readers, were presented in white Arial font, on a black background. Words in each pair differed by one letter (for example, “tack” vs. “tank”). The second letter differed in 14 pairs, and the third letter differed in 15. As shown in Fig. 5c, participants viewed a word centered over fixation for 750 ms (long enough for encoding by children). After a 150 ms interval with just a fixation cross, participants saw a second word centered over fixation for 150 ms. A randomly jittered 1500 to 2500 ms interval separated trials. Participants indicated, via key press, whether the two stimuli were the same or different, for 96 trials (half same, half different). Participants completed 12 practice trials and received feedback, and no direct feedback was provided during the task.

### Quantification and Statistical Analysis

#### Quantification

Data preprocessing steps were adapted from prior fMRI work with this patient population^22^. Anatomical data were defaced with PyDeface^64^, and then, using FreeSurfer (v7.1.0), underwent motion correction^65^, intensity normalization^66^, and skull stripping^67^. All fMRI preprocessing and derivation of dependent measures were then completed with Analysis of Functional Neuroimages^68,69^. EPI data, which were acquired in oblique orientation, were transformed to a cardinal grid. Volumes were slice-time-corrected. The volume with least motion was aligned to the anatomical data with a local Pearson correlation cost function^70^. Data were scaled to a mean of 100 per run and were not spatially smoothed to preserve resolution.

Volumes were discarded if the Euclidean norm of motion derivatives exceeded 1 cm or if outlier values (approximately 5.5 median absolute deviations away from the median) were detected in more than 10% of voxels. To optimize registration of brains with complete hemispheric surgery (e.g., minimal residual tissue of the operated hemisphere) in AFNI, after defacement and prior to preprocessing, anatomical data were mirrored^71^. Additionally, only EPI data from the preserved hemisphere were analyzed for all patients.

The first two runs of the category localizer data, which were used to define ROIs, were concatenated and preprocessed independently from the third run, which was held out to independently derive CS amplitudes. Separately for each the concatenated first two runs and the third run, a GLM was fit with each visual category (faces, objects, words, houses, and scrambled objects) as predictors of a boxcar function. Regressors for motion in each of the six coordinate directions (translation: x, y, and z; and rotation: pitch, roll, yaw) and scanner drift were also modeled. Temporal autocorrelations were regressed out of the data. To restrict analyses to only functionally relevant parcels, each participant’s brain was aligned to the Human Connectome Project Multi-Modal Parcellation (v1) atlas^41^ in Montreal Neurological Institute space and then back-transformed into native space^22,72^. A mask was created, separately in the LH and RH, that only included voxels within the ventral stream visual cortex parcel^41^. To correct for multiple comparisons, within this mask for each hemisphere, Monte Carlo simulations of noise data sets (thresholded at *p* < 0.001) were run 10,000 times to approximate the minimum size of a cluster of significant voxels that would not be expected due to chance alone (at an α of 0.05)^73,74^. With the first two runs of the data, a CS ROI was then defined, as the collection of clustered positive voxels for the balanced contrast of the β-weights for the given stimulus category against all other categories, above the designated cluster threshold and within the ventral stream visual cortex mask. CS amplitude was defined as the median β-weight across all voxels for a given stimulus category on the third run of data, within the corresponding CS ROI (defined with the first two runs of the data).

fMRI data from the adaptation paradigm were preprocessed as above, except all six runs were concatenated, and a Gamma basis function instead of a boxcar, was convolved with the hemodynamic response function, given the abbreviated block lengths. A GLM was fit to the basis function with each block type (e.g., same/alternating/different x faces/objects/words) as predictors. The median β-weight across every voxel was calculated for each block condition within the preferred CS-ROI (e.g., same faces in the face-selective ROI, different words in the word-selective ROI).

For the behavioral data, mean accuracy was computed per participant on each task. Mean accuracy on a given task was discarded for an individual participant if it was at or below 33% on the face and object recognition tasks (with three response choices) or 50% on the word recognition task (with two response choices). No participants performed below chance on either the face or object recognition tasks. One control patient performed at chance on the word recognition task.

#### Statistical Analysis

For each dependent measure (e.g., CS amplitude, RS, behavioral accuracy), the same statistical approach was applied. First, following the procedures outlined by Fox & Weisberg^35^ and Brown^34^, a LMEM was fit with the complete interaction of relevant predictors (e.g., group, stimulus category, hemisphere) as a fixed effect, mean-centered age as a fixed covariate, and participant modeled as a random intercept. The LRT was used to compare the model with the complete interaction vs. each lower-order model without the complete interaction (e.g. a model with a three-way interaction was compared to a model with all two-way interactions, each model with a two-way interaction was compared to a model without each two-way interaction, etc.; see Tables in Supplemental Information for listing of all factors in a model and then the selection of the best fit). Models with the highest order interaction were compared to the next most complex model, and so on. In these stepwise comparisons, if two models provided significantly different fits of the data, the one with the lower AIC was tested, and no further comparisons were made. All models included group (the key predictor of interest) and age (the primary covariate, expected to affect each dependent variable^12,30,75^). In approximating fixed effects, the maximum log-likelihood criterion was optimized^76^, and the Bound Optimization by Quadratic Approximation algorithm was applied^77^. Wald χ^2^-tests were employed to calculate *p*-values for each effect^35^. If an interaction effect was statistically significant, post hoc contrasts on the EMMs for meaningful factor comparisons were calculated^78^.

Two statistical approaches were adopted from prior work with small (though smaller) sample sizes of this patient population^20–22^. First, the three groups (OTC patients, control patients, and TD controls) were compared to one another using permutation testing. For each model evaluated, group labels were shuffled 1000 times without replacement, and the model was refit. A Wald χ^2^-test was run on each refitted model, and a *p*-value was calculated as the proportion of instances across the permutations in which the derived χ^2^-value on a refitted model was greater than the true χ^2^-value. In addition, as group is a three-level variable, if the group effect was statistically significant, post hoc contrast testing was performed. On each of the shuffled datasets, a *z*-score was computed for each pairwise comparison, and the proportion of instances across the permutations in which each *z*-score was greater than each true *z*-score was computed. Additionally, to further address the small sample size, an approach developed by Crawford et al.^79^ to compare individual patient measures to small control samples (while regressing out age) was adopted. Individual comparisons between patients and TD controls were performed separately with data from TD controls’ LHs and RHs.

Finally, RSA was conducted to examine the pattern of activation in the ventral stream. For each participant, an activation RDM was constructed by computing the similarity between each block type (e.g., same/alternating/different x faces/objects/words) across all voxels from the ventral stream visual cortex parcel using Pearson correlation. Three model RDMs were then constructed based on properties of the ventral stream related to CS and RS. Each participant’s activation RDM was compared to each of the three model RDMs using Spearman’s correlation. Spearman’s correlation maps for each participant were Fisher transformed. The effects of group, hemisphere, and model type (CS, RS, CS-RS) on the Fisher transformation was analyzed using the same approach as for predicting other dependent measures (e.g., CS magnitude, behavioral accuracy) as described above.

All statistical analyses were performed in R^80^, with the exception of the RSA, which was conducted in MATLAB. For a complete list of R packages used, see Table S17. Select code was written with the assistance of ChatGPT. α criterion for statistical significance was 0.05. A BF was also calculated by comparing a model with a given effect to a null model without that effect: BFs above 3 or below 0.33 can be interpreted as evidence supporting the null or alternative hypothesis, respectively^36,81^. The Benjamini-Hochberg correction^37^ was applied to *p*-values of each set of post hoc comparisons and each set of Crawford comparisons (e.g., all patients compared to one hemisphere of the TD controls on a given measure).

## Acknowledgements

This research was supported by Award Numbers T32GM008208 and T32GM081760 from the National Institute of General Medical Sciences, fellowship #847566 from the American Epilepsy Society, a scholarship from the American Psychological Foundation, a scholarship from the University of Pittsburgh MD-PhD program to MCG; a fellowship under Grant #DGE2140739 from the National Science Foundation to SR; and by R01EY027018 from the National Eye Institute to MB and CP. MB also acknowledges support from P30 CORE Award (EY08098) from the National Eye Institute, NIH, and unrestricted supporting funds from The Research to Prevent Blindness Inc, NY, and the Eye & Ear Foundation of Pittsburgh. The content is solely the responsibility of the authors and does not necessarily represent the official views of NIGMS, AES, APF, NSF, or NEI. We are grateful to the Pediatric Epilepsy Surgery Alliance and its Chief Executive Officer Monika Jones, JD for assistance with participant recruitment. We also thank Scott Kurdilla, Mark Vignone, and Debbie Viszlay for assistance with data acquisition; and Drs. Vladislav Ayzenberg, Nicholas Blauch, Carl Olson, David Plaut, Michael Tarr, and Timothy Verstynen for their helpful discussions and feedback. Face images for category localizer courtesy of Michael J. Tarr, Carnegie Mellon University, http://www.tarrlab.org/; funding provided by NSF award 0339122. Greeble stimuli courtesy of Suzy Scherf. We especially thank the participants and their families for their time and cooperation.

## Author Contributions

**Conceptualization**: MCG, EF, MB. **Methodology**: MCG, AMSM, SL, SR, EF, MB. **Software**: MCG, AMSM, SL. **Validation**: MCG, SL. **Formal analysis**: MCG, SL. **Investigation**: MCG, AMSM, SR. **Data curation**: MCG, AMSM. **Resources**: CP. **Writing - Original Draft**: MCG. **Writing - Review & Editing**: AMSM, SL, SR, EF, CP, MB. **Visualization**: MCG, SL. **Supervision**: MB. **Project administration**: MCG, AMSM, MB. **Funding acquisition**: MCG, CP, MB.

## References

1. Grill-Spector, K. & Malach, R. The human visual cortex. Annu. Rev. Neurosci. 27, 649–677 (2004).

2. Grill-Spector, K. & Weiner, K. S. The functional architecture of the ventral temporal cortex and its role in categorization. Nat. Rev. Neurosci. 15, 536–548 (2014).

3. Barttfeld, P. et al. A lateral-to-mesial organization of human ventral visual cortex at birth. Brain Struct. Funct. 223, 3107–3119 (2018).

4. Boring, M. J. et al. Multiple adjoining word- and face-selective regions in ventral temporal cortex exhibit distinct dynamics. J. Neurosci. 41, 6314–6327 (2021).

5. Matsuo, T. et al. Alternating zones selective to faces and written words in the human ventral occipitotemporal cortex. Cereb. Cortex 25, 1265–1277 (2015).

6. Behrmann, M. & Plaut, D. C. Hemispheric organization for visual object recognition: A theoretical account and empirical evidence. Perception 49, 373–404 (2020).

7. Bourne, J. A. Unravelling the development of the visual cortex: implications for plasticity and repair. J. Anat. 217, 449–468 (2010).

8. Stiles, J., Reilly, J., Paul, B. & Moses, P. Cognitive development following early brain injury: evidence for neural adaptation. Trends Cogn. Sci. (Regul. Ed.) 9, 136–143 (2005).

9. Germine, L. T., Duchaine, B. & Nakayama, K. Where cognitive development and aging meet: face learning ability peaks after age 30. Cognition 118, 201–210 (2011).

10. Scherf, K. S., Behrmann, M., Humphreys, K. & Luna, B. Visual category-selectivity for faces, places and objects emerges along different developmental trajectories. Dev. Sci. 10, F15–30 (2007).

11. Haist, F., Adamo, M., Han Wazny, J., Lee, K. & Stiles, J. The functional architecture for face-processing expertise: FMRI evidence of the developmental trajectory of the core and the extended face systems. Neuropsychologia 51, 2893–2908 (2013).

12. Nordt, M. et al. Cortical recycling in high-level visual cortex during childhood development. *Nat*. Hum. Behav. 5, 1686–1697 (2021).

13. Dehaene-Lambertz, G., Monzalvo, K. & Dehaene, S. The emergence of the visual word form: Longitudinal evolution of category-specific ventral visual areas during reading acquisition. PLoS Biol. 16, e2004103 (2018).

14. Feng, X., Monzalvo, K., Dehaene, S. & Dehaene-Lambertz, G. Evolution of reading and face circuits during the first three years of reading acquisition. Neuroimage 259, 119394 (2022).

15. Yeatman, J. D. et al. Reading instruction causes changes in category-selective visual cortex. Brain Res. Bull. 110958 (2024). doi:10.1016/j.brainresbull.2024.110958

16. Gomez, J., Barnett, M. & Grill-Spector, K. Extensive childhood experience with Pokémon suggests eccentricity drives organization of visual cortex. *Nat*. Hum. Behav. 3, 611–624 (2019).

17. Behrmann, M. & Plaut, D. C. A vision of graded hemispheric specialization. Ann. N. Y. Acad. Sci. 1359, 30–46 (2015).

18. Kolb, B., Mychasiuk, R., Muhammad, A. & Gibb, R. Brain plasticity in the developing brain. Prog. Brain Res. 207, 35–64 (2013).

19. Kolb, B. & Gibb, R. Brain plasticity and behaviour in the developing brain. J Can Acad Child Adolesc Psychiatry 20, 265–276 (2011).

20. Liu, T. T. et al. Successful Reorganization of Category-Selective Visual Cortex following Occipito-temporal Lobectomy in Childhood. Cell Rep. 24, 1113–1122.e6 (2018).

21. Liu, T. T., Freud, E., Patterson, C. & Behrmann, M. Perceptual Function and Category-Selective Neural Organization in Children with Resections of Visual Cortex. J. Neurosci. 39, 6299–6314 (2019).

22. Maallo, A. M. S. et al. Large-scale resculpting of cortical circuits in children after surgical resection. Sci. Rep. 10, 21589 (2020).

23. Kriegeskorte, N., Mur, M. & Bandettini, P. Representational similarity analysis - connecting the branches of systems neuroscience. Front. Syst. Neurosci. 2, 4 (2008).

24. Barron, H. C., Garvert, M. M. & Behrens, T. E. J. Repetition suppression: a means to index neural representations using BOLD? *Philos. Trans. R. Soc. Lond. B*, Biol. Sci. 371, (2016).

25. Grill-Spector, K. & Malach, R. fMR-adaptation: a tool for studying the functional properties of human cortical neurons. Acta Psychol. (Amst.) 107, 293–321 (2001).

26. Grill-Spector, K., Henson, R. & Martin, A. Repetition and the brain: neural models of stimulus-specific effects. Trends Cogn. Sci. (Regul. Ed.) 10, 14–23 (2006).

27. Malach, R. Targeting the functional properties of cortical neurons using fMR-adaptation. Neuroimage 62, 1163–1169 (2012).

28. Konen, C. S. & Kastner, S. Two hierarchically organized neural systems for object information in human visual cortex. Nat. Neurosci. 11, 224–231 (2008).

29. Scherf, K. S., Luna, B., Avidan, G. & Behrmann, M. “What” precedes “which”: developmental neural tuning in face- and place-related cortex. Cereb. Cortex 21, 1963– 1980 (2011).

30. Granovetter, M. C., Robert, S., Ettensohn, L. & Behrmann, M. With childhood hemispherectomy, one hemisphere can support-but is suboptimal for-word and face recognition. Proc. Natl. Acad. Sci. USA 119, e2212936119 (2022).

31. Simmons, C. et al. Holistic processing and face expertise after pediatric resection of occipitotemporal cortex. Neuropsychologia 194, 108789 (2024).

32. Pinabiaux, C., Save-Pédebos, J., Dorfmüller, G., Jambaqué, I. & Bulteau, C. The hidden face of hemispherectomy: Visuo-spatial and visuo-perceptive processing after left or right functional hemispherectomy in 40 children. Epilepsy Behav. 134, 108821 (2022).

33. Commission on Neurosurgery of the International League Against Epilepsy (ILAE) 1997- 2001: et al. Proposal for a New Classification of Outcome with Respect to Epileptic Seizures Following Epilepsy Surgery. Epilepsia 42, 282–286 (2008).

34. Brown, V. A. An introduction to linear mixed-effects modeling in R. Advances in Methods and Practices in Psychological Science 4, 251524592096035 (2021).

35. Fox, J. & Weisberg, S. An {R} Companion to Applied Regression. (Sage, 2019).

36. Wagenmakers, E.-J. A practical solution to the pervasive problems of p values. Psychon. Bull. Rev. 14, 779–804 (2007).

37. Benjamini, Y. & Yekutieli, D. The control of the false discovery rate in multiple testing under dependency. Ann. Statist. 29, 1165–1188 (2001).

38. Rosenthal, G. et al. Altered topology of neural circuits in congenital prosopagnosia. Elife 6, (2017).

39. Freud, E. et al. Three-Dimensional Representations of Objects in Dorsal Cortex are Dissociable from Those in Ventral Cortex. Cereb. Cortex 27, 422–434 (2017).

40. Crawford, J. R. & Garthwaite, P. H. Single-case research in neuropsychology: a comparison of five forms of t-test for comparing a case to controls. Cortex 48, 1009–1016 (2012).

41. Glasser, M. F. et al. A multi-modal parcellation of human cerebral cortex. Nature 536, 171– 178 (2016).

42. Righi, G., Peissig, J. J. & Tarr, M. J. Recognizing disguised faces. Vis. cogn. 20, 143–169 (2012).

43. Lundqvist, D., Flykt, A. & Öhman, A. The Karolinska Directed Emotional Faces - KDEF. (CD ROM from Department of Clinical Neuroscience, Psychology section, Karolinska Institutet, ISBN 91-630-7164-9, 1998).

44. Croydon, A., Pimperton, H., Ewing, L., Duchaine, B. C. & Pellicano, E. The Cambridge Face Memory Test for Children (CFMT-C): a new tool for measuring face recognition skills in childhood. Neuropsychologia 62, 60–67 (2014).

45. Duchaine, B. & Nakayama, K. The Cambridge Face Memory Test: results for neurologically intact individuals and an investigation of its validity using inverted face stimuli and prosopagnosic participants. Neuropsychologia 44, 576–585 (2006).

46. Dalrymple, K. A., Garrido, L. & Duchaine, B. Dissociation between face perception and face memory in adults, but not children, with developmental prosopagnosia. Dev Cogn Neurosci 10, 10–20 (2014).

47. Dalrymple, K. A., Elison, J. T. & Duchaine, B. Face-specific and domain-general visual processing deficits in children with developmental prosopagnosia. Q J Exp Psychol (Colchester) 70, 259–275 (2017).

48. Dundas, E. M., Plaut, D. C. & Behrmann, M. The joint development of hemispheric lateralization for words and faces. J. Exp. Psychol. Gen. 142, 348–358 (2013).

49. Behrmann, M. & Plaut, D. C. Bilateral hemispheric processing of words and faces: evidence from word impairments in prosopagnosia and face impairments in pure alexia. Cereb. Cortex 24, 1102–1118 (2014).

50. Landis, T., Cummings, J. L., Christen, L., Bogen, J. E. & Imhof, H. G. Are unilateral right posterior cerebral lesions sufficient to cause prosopagnosia? Clinical and radiological findings in six additional patients. Cortex 22, 243–252 (1986).

51. Damasio, A. R. & Damasio, H. The anatomic basis of pure alexia. Neurology 33, 1573– 1583 (1983).

52. Bate, S. & Bennetts, R. J. The rehabilitation of face recognition impairments: a critical review and future directions. Front. Hum. Neurosci. 8, 491 (2014).

53. Starrfelt, R., Olafsdóttir, R. R. & Arendt, I.-M. Rehabilitation of pure alexia: a review. Neuropsychol Rehabil 23, 755–779 (2013).

54. de Schonen, S., Mancini, J., Camps, R., Maes, E. & Laurent, A. Early brain lesions and face-processing development. Dev. Psychobiol. 46, 184–208 (2005).

55. Albonico, A. & Barton, J. Progress in perceptual research: the case of prosopagnosia. [version 1; peer review: 2 approved]. F1000Res. 8, (2019).

56. Olulade, O. A. et al. The neural basis of language development: Changes in lateralization over age. Proc. Natl. Acad. Sci. USA 117, 23477–23483 (2020).

57. Robert, S., Granovetter, M. C., Patterson, C. & Behrmann, M. Investigation of hemispheric functional organization after pediatric epilepsy surgery with naturalistic neuroimaging. J. Vis. 22, 4454 (2022).

58. Robert, S., Granovetter, M. C., Patterson, C. & Behrmann, M. Hemispheric functional organization, as revealed by naturalistic neuroimaging, in pediatric epilepsy patients with cortical resections.

59. Ayzenberg, V., Granovetter, M. C., Robert, S., Patterson, C. & Behrmann, M. Differential functional reorganization of ventral and dorsal visual pathways following childhood hemispherectomy. Dev Cogn Neurosci 64, 101323 (2023).

60. Eriksson, M. H. et al. Long-term neuropsychological trajectories in children with epilepsy: does surgery halt decline? Brain (2024). doi:10.1093/brain/awae121

61. Brainard, D. H. The Psychophysics Toolbox. Spat Vis 10, 433–436 (1997).

62. Stigliani, A., Weiner, K. S. & Grill-Spector, K. Temporal Processing Capacity in High-Level Visual Cortex Is Domain Specific. J. Neurosci. 35, 12412–12424 (2015).

63. Gauthier, I., Williams, P., Tarr, M. J. & Tanaka, J. Training “greeble” experts: a framework for studying expert object recognition processes. Vision Res. 38, 2401–2428 (1998).

64. Gulban, O. F., et al. PyDeface. (2019).

65. Reuter, M., Rosas, H. D. & Fischl, B. Highly accurate inverse consistent registration: a robust approach. Neuroimage 53, 1181–1196 (2010).

66. Sled, J. G., Zijdenbos, A. P. & Evans, A. C. A nonparametric method for automatic correction of intensity nonuniformity in MRI data. IEEE Trans. Med. Imaging 17, 87–97 (1998).

67. Ségonne, F. et al. A hybrid approach to the skull stripping problem in MRI. Neuroimage 22, 1060–1075 (2004).

68. Cox, R. W. AFNI: software for analysis and visualization of functional magnetic resonance neuroimages. Comput. Biomed. Res. 29, 162–173 (1996).

69. Cox, R. W. & Hyde, J. S. Software tools for analysis and visualization of fMRI data. NMR Biomed. 10, 171–178 (1997).

70. Saad, Z. S. et al. A new method for improving functional-to-structural MRI alignment using local Pearson correlation. Neuroimage 44, 839–848 (2009).

71. Granovetter, M. C., Maallo, A. M. S., Patterson, C., Glen, D. & Behrmann, M. Morphometrics of the preserved post-surgical hemisphere in paediatric drug-resistant epilepsy. BioRxiv (2023). doi:10.1101/2023.09.24.559189

72. Glen, D., Levenstein, J., Granovetter, M., Maallo, A. M. S. & Behrmann, M. LARGE Lesion Brain Alignment with AFNI. (2021).

73. Cox, R. W., Chen, G., Glen, D. R., Reynolds, R. C. & Taylor, P. A. fMRI clustering and false-positive rates. Proc. Natl. Acad. Sci. USA 114, E3370–E3371 (2017).

74. Cox, R. W., Chen, G., Glen, D. R., Reynolds, R. C. & Taylor, P. A. FMRI Clustering in AFNI: False-Positive Rates Redux. Brain Connect. 7, 152–171 (2017).

75. Nordt, M., Hoehl, S. & Weigelt, S. The use of repetition suppression paradigms in developmental cognitive neuroscience. Cortex 80, 61–75 (2016).

76. Bates, D., Mächler, M., Bolker, B. & Walker, S. Fitting linear mixed-effects models using lme4. J. Stat. Softw. 67, 1–48 (2015).

78. Powell, M. J. D. The BOBYQA Algorithm for Bound Constrained Optimization without Derivatives. (2009).

78. Lenth, R. V. emmeans: Estimated Marginal Means, aka Least-Squares Means. (2021).

79. Crawford, J. R., Garthwaite, P. H. & Ryan, K. Comparing a single case to a control sample: testing for neuropsychological deficits and dissociations in the presence of covariates. Cortex 47, 1166–1178 (2011).

80. R Core Team. R: A Language and Environment for Statistical Computing. (R Foundation for Statistical Computing, 2020).

81. Lee, M. D. & Wagenmakers, E.-J. Bayesian cognitive modeling: A practical course. (Cambridge University Press, 2013). doi:10.1017/CBO9781139087759

