## Supplementary Information for "Functional Resilience of the Neural Visual Recognition System Post-Pediatric Occipitotemporal Resection"

### Supplemental Information

**Table S1**: *Model Selection (Category Selectivity)*

| Models | | | |
|---|---|---|---|
| # | Model | | AIC |
| 1 | CS ~ group * hemi * stim + age + (1\|ID) | | 326.66 |
| 2 | CS ~ group * hemi + group * stim + hemi * stim + age + (1\|ID) | | 322.14 |
| 3 | CS ~ group * hemi + group * stim + age + (1\|ID) | | 320.79 |
| 4 | CS ~ group * hemi + hemi * stim + age + (1\|ID) | | 316.95 |
| 5 | CS ~ group * stim + hemi * stim + age + (1\|ID) | | 320.84 |
| 6 | CS ~ hemi * stim + group + age + (1\|ID) | | 315.73 |
| 7 | CS ~ group * hemi + stim + age + (1\|ID) | | 315.11 |
| 8 | CS ~ group * stim + hemi + age + (1\|ID) | | 319.59 |
| 9 | **CS ~ group + stim + hemi + age + (1\|ID)** | | 313.98 |
| 10 | CS ~ group + hemi + age + (1\|ID) | | 316.83 |
| 11 | CS ~ group + stim + age + (1\|ID) | | 311.98 |
| LRTs | | | |
| # vs. # | $\chi^2$ | Df | *p* |
| 1 vs. 2 | 3.48 | 4 | 0.48 |
| 2 vs. 3 | 2.65 | 2 | 0.27 |
| 2 vs. 4 | 2.81 | 4 | 0.59 |
| 2 vs. 5 | 2.70 | 2 | 0.26 |
| 3 vs. 4 | 0.16 | 2 | 0.92 |
| 4 vs. 5 | 0.11 | 2 | 0.95 |
| 3 vs. 7 | 2.32 | 4 | 0.68 |
| 3 vs. 8 | 2.80 | 2 | 0.25 |
| 4 vs. 6 | 2.78 | 2 | 0.25 |
| 4 vs. 7 | 2.16 | 2 | 0.34 |
| 5 vs. 6 | 2.90 | 4 | 0.58 |
| 5 vs. 8 | 2.75 | 2 | 0.25 |
| 6 vs. 8 | 0.14 | 2 | 0.93 |
| 7 vs. 8 | 0 | 2 | ~ 1 |
| 6 vs. 9 | 2.25 | 2 | 0.33 |
| 7 vs. 9 | 2.87 | 2 | 0.24 |
| 8 vs. 9 | 2.39 | 4 | 0.66 |
| 9 vs. 10 | 6.85 | 2 | **0.03** |
| 9 vs. 11 | < 0.01 | 1 | 0.97 |

Selected model and significant *p*-value are bolded. LRT = likelihood ratio test. AIC = Akaike information criterion. Df = degrees of freedom. CS = category selectivity amplitude. hemi = hemisphere. stim = stimulus category.

**Table S2**: *Post Hoc Comparisons (Category Selectivity ~ Stimulus Category)*

| Comparison | *z* | *p* |
|---|---|---|
| Faces - Objects | 0.51 | 0.61 |
| Faces - Words | 2.55 | **0.03** |
| Objects - Words | 2.07 | 0.06 |

Significant *p*-value is bolded.

**Table S3**: *Crawford Statistics (Category Selectivity, Repetition Suppression, Behavior)*

| ID | Stimulus Category | *p* (comparison | *p* (comparison | *p* (comparison | *p* (comparison | *p* (comparison |
|---|---|---|---|---|---|---|

| | | to TD LH on CS) | to TD RH on CS) | to TD LH on RS) | to TD RH on RS) | to TD on behavior) |
|---|---|---|---|---|---|---|
| Preserved LH OTC Patients | | | | | | |
| sub-004 | Faces | 0.84 | 0.84 | 0.86 | 0.85 | 0.74 |
| | Objects | 0.82 | 0.84 | 0.83 | 0.80 | 0.75 |
| | Words | - | - | - | - | 0.84 |
| sub-077 | Faces | 0.84 | 0.84 | 0.82 | 0.85 | 0.63 |
| | Objects | 0.82 | 0.84 | 0.20 | 0.80 | 0.30 |
| | Words | 0.79 | 0.83 | 0.85 | 0.79 | 0.49 |
| sub-079 | Faces | 0.84 | 0.84 | 0.86 | 0.85 | 0.74 |
| | Objects | 0.82 | 0.84 | 0.83 | 0.80 | 0.62 |
| | Words | 0.79 | 0.83 | 0.85 | 0.79 | 0.84 |
| sub-089 | Faces | 0.84 | 0.84 | 0.52 | 0.70 | **0.01** |
| | Objects | 0.82 | 0.84 | 0.83 | 0.80 | 0.27 |
| | Words | 0.83 | 0.83 | 0.85 | 0.79 | 0.84 |
| sub-091 | Faces | 0.84 | 0.84 | 0.53 | 0.85 | 0.63 |
| | Objects | 0.82 | 0.84 | 0.83 | 0.80 | 0.15 |
| | Words | 0.83 | 0.83 | 0.85 | 0.79 | 0.84 |
| Preserved RH OTC Patients | | | | | | |
| sub-066 | Faces | 0.84 | 0.84 | 0.42 | 0.70 | **0.03** |
| | Objects | 0.82 | 0.84 | 0.86 | 0.80 | - |
| | Words | 0.83 | 0.83 | 0.85 | 0.79 | 0.49 |
| sub-069 | Faces | - | - | - | - | **0.01** |
| | Objects | 0.82 | 0.84 | 0.83 | 0.80 | - |
| | Words | 0.63 | 0.83 | 0.85 | 0.79 | **0.01** |
| sub-090 | Faces | 0.84 | 0.84 | 0.86 | 0.85 | 0.74 |
| | Objects | 0.82 | 0.84 | 0.83 | 0.80 | 0.20 |
| | Words | 0.79 | 0.83 | 0.85 | 0.79 | 0.84 |
| sub-092 | Faces | 0.84 | 0.84 | 0.86 | 0.85 | 0.07 |
| | Objects | 0.82 | 0.84 | 0.83 | 0.80 | 0.41 |
| | Words | 0.63 | 0.83 | 0.85 | 0.79 | 0.84 |
| Preserved LH Control Patients | | | | | | |
| sub-072 | Faces | 0.84 | 0.84 | 0.86 | 0.85 | 0.74 |
| | Objects | 0.82 | 0.84 | 0.83 | 0.80 | 0.20 |
| | Words | 0.63 | 0.83 | 0.85 | 0.79 | 0.84 |
| sub-073 | Faces | 0.84 | 0.84 | 0.86 | 0.85 | 0.74 |
| | Objects | 0.82 | 0.84 | 0.83 | 0.80 | 0.75 |
| | Words | 0.83 | 0.83 | 0.85 | 0.79 | 0.84 |

| | | | | | | |
|---|---|---|---|---|---|---|
| sub-075 | Faces | 0.84 | 0.84 | 0.53 | 0.70 | **0.01** |
| | Objects | 0.82 | 0.84 | 0.58 | 0.80 | 0.20 |
| | Words | - | - | - | - | - |
| sub-080 | Faces | 0.84 | 0.84 | 0.86 | 0.85 | 0.74 |
| | Objects | 0.82 | 0.84 | 0.83 | 0.80 | 0.41 |
| | Words | - | - | - | - | 0.84 |
| sub-086 | Faces | 0.84 | 0.84 | 0.86 | 0.85 | 0.84 |
| | Objects | 0.82 | 0.84 | 0.83 | 0.80 | 0.71 |
| | Words | 0.63 | 0.83 | 0.69 | 0.32 | 0.84 |
| Preserved RH Control Patients | | | | | | |
| sub-007 | Faces | 0.84 | 0.84 | 0.86 | 0.85 | 0.74 |
| | Objects | 0.82 | 0.84 | 0.83 | 0.80 | 0.75 |
| | Words | - | - | - | - | 0.84 |
| sub-045 | Faces | 0.84 | 0.84 | 0.86 | 0.85 | 0.63 |
| | Objects | 0.82 | 0.84 | 0.83 | 0.80 | 0.75 |
| | Words | - | - | - | - | 0.84 |
| sub-070 | Faces | 0.84 | 0.84 | 0.53 | 0.85 | 0.74 |
| | Objects | 0.82 | 0.84 | 0.83 | 0.80 | 0.58 |
| | Words | 0.63 | 0.83 | 0.85 | 0.79 | 0.84 |
| sub-076 | Faces | 0.84 | 0.84 | 0.42 | 0.70 | 0.74 |
| | Objects | - | - | - | - | 0.58 |
| | Words | - | - | - | - | 0.84 |
| sub-078 | Faces | 0.84 | 0.84 | **0.01** | 0.11 | 0.74 |
| | Objects | 0.82 | 0.84 | 0.83 | 0.80 | 0.51 |
| | Words | 0.83 | 0.83 | 0.85 | 0.79 | 0.84 |
| sub-081 | Faces | 0.84 | 0.84 | 0.86 | 0.85 | 0.63 |
| | Objects | 0.82 | 0.84 | 0.83 | 0.80 | 0.58 |
| | Words | 0.83 | 0.83 | 0.85 | 0.79 | 0.84 |
| sub-082 | Faces | 0.84 | 0.84 | 0.86 | 0.85 | 0.74 |
| | Objects | 0.82 | 0.84 | 0.83 | 0.80 | 0.58 |
| | Words | 0.83 | 0.83 | 0.85 | 0.79 | 0.84 |

Significant *p*-values are bolded. TD = typically developing controls. LH = left hemisphere. RH = right hemisphere. CS = category selectivity amplitude. RS = repetition suppression magnitude.

**Table S4**: *Model Selection (Responsivity to Adaptation Experiment)*

| Models | | |
|---|---|---|
| # | Model | AIC |
| 1 | resp ~ group * cond * hemi * stim + age + (1\|ID) | 445.17 |
| 2 | **resp ~ group * cond * hemi + group * cond * stim + group * hemi * stim + cond * hemi * stim + age + (1\|ID)** | 430.30 |
| 3 | resp ~ group * cond * stim + group * hemi * stim + cond * hemi * stim + age + (1\|ID) | 423.17 |

| | | |
|---|---|---|
| 4 | resp ~ group * cond * hemi + group * hemi * stim + cond * hemi * stim + age + (1\|ID) | 416.86 |
| 5 | resp ~ group * cond * hemi + group * cond * stim + cond * hemi * stim + age + (1\|ID) | 427.31 |
| 6 | resp~ group * cond * hemi + group * cond * stim + group * hemi * stim + age + (1\|ID) | 423.59 |
| 7 | resp ~ group * cond * hemi + group * cond * stim + age + (1\|ID) | 428.72 |
| 8 | resp ~ group * cond * hemi + group * hemi * stim + age + (1\|ID) | 403.18 |
| 9 | resp ~ group * cond * hemi + cond * hemi * stim + age + (1\|ID) | 416.86 |
| 10 | resp ~ group * cond * stim + group * hemi * stim + age + (1\|ID) | 412.54 |
| 11 | resp ~ group * cond * stim + cond * hemi * stim + age + (1\|ID) | 420.72 |
| 12 | resp ~ group * hemi * stim + cond * hemi * stim + age + (1\|ID) | 401.93 |

LRTs

| # vs. # | $\chi^2$ | Df | *p* |
|---|---|---|---|
| 1 vs. 2 | 1.13 | 8 | ~ 1 |
| 2 vs. 3 | 0.87 | 4 | 0.93 |
| 2 vs. 4 | 2.56 | 8 | 0.96 |
| 2 vs. 5 | 5.01 | 4 | 0.29 |
| 2 vs. 6 | 1.29 | 4 | 0.86 |
| 3 vs. 4 | 1.69 | 4 | 0.79 |
| 4 vs. 5 | 0 | 4 | ~ 1 |
| 4 vs. 6 | 1.27 | 4 | 0.87 |
| 3 vs. 10 | 1.37 | 6 | 0.97 |
| 3 vs. 11 | 9.55 | 6 | 0.14 |
| 3 vs. 12 | 2.76 | 12 | ~ 1 |
| 4 vs. 8 | 2.32 | 8 | 0.97 |
| 4 vs. 9 | 15.41 | 8 | 0.05 |
| 4 vs. 12 | 1.07 | 8 | ~ 1 |
| 5 vs. 7 | 13.41 | 6 | **0.04** |
| 5 vs. 9 | 12.97 | 12 | 0.37 |
| 5 vs. 11 | 5.41 | 6 | 0.49 |
| 6 vs. 7 | 17.13 | 6 | **0.01** |
| 6 vs. 8 | 3.59 | 12 | 0.99 |
| 6 vs. 10 | 0.94 | 6 | 0.99 |

Selected model and significant *p*-value are bolded. Note that since models 5 and 6 both better fit the data than model 7 with fewer parameters, and since model 2 is the model with the fewest parameters that nonetheless contains all the parameters in models 5 and 6, model 2 was selected. LRT = likelihood ratio test. AIC = Akaike information criterion. Df = degrees of freedom. resp = responsivity. cond = condition (same, alternating, different). hemi = hemisphere. stim = stimulus category.

**Table S5**: *Post Hoc Comparisons (Responsivity ~ Hemisphere x Stimulus Category)*

| Comparison | *z* | *p* |
|---|---|---|
| LH Faces - RH Faces | 0.14 | 0.99 |
| LH Faces - LH Objects | 3.71 | **< 0.001** |
| LH Faces - LH Words | -4.16 | **< 0.001** |
| RH Faces - RH Objects | 4.92 | **< 0.001** |
| RH Faces - RH Words | 0.02 | 0.99 |
| LH Objects - RH Objects | 1.15 | 0.32 |
| LH Objects - LH Words | -7.42 | **< 0.001** |
| RH Objects - RH Words | -3.81 | **< 0.001** |
| LH Words - RH Words | 2.89 | **0.01** |

Significant *p*-values are bolded. LH = left hemisphere. RH = right hemisphere.

**Table S6**: *Post Hoc Comparisons (Responsivity ~ Group x Stimulus Category)*

| Stimulus Category | Comparison | *z* | *p* | *p* (permutation testing) |
|---|---|---|---|---|
| Faces | OTC - TD | -0.13 | 0.96 | 0.95 |
| | OTC - CP | -0.08 | 0.96 | 0.95 |
| | TD - CP | 0.05 | 0.96 | 0.95 |
| Objects | OTC - TD | -0.59 | 0.96 | 0.95 |
| | OTC - CP | 0.36 | 0.96 | 0.95 |
| | TD - CP | 1.09 | 0.83 | 0.79 |
| Words | OTC - TD | -0.54 | 0.96 | 0.95 |
| | OTC - CP | -1.96 | 0.30 | 0.79 |
| | TD - CP | -1.84 | 0.30 | 0.79 |

OTC = occipitotemporal cortex patients. TD = typically developing controls. CP = control patients.

**Table S7**: *Crawford Statistics (Responsivity to Adaptation Experiment)*

| ID | Stimulus Category | Condition | *p* (comparison to TD LH) | *p* (comparison to TD RH) |
|---|---|---|---|---|
| Preserved LH OTC Patients | | | | |
| sub-004 | Faces | Same | 0.91 | 0.90 |
| | | Alt | 0.91 | 0.90 |
| | | Diff | 0.91 | 0.90 |
| | Objects | Same | 0.88 | 0.91 |
| | | Alt | 0.88 | 0.91 |
| | | Diff | 0.88 | 0.91 |
| | Words | Same | - | - |
| | | Alt | - | - |
| | | Diff | - | - |
| sub-077 | Faces | Same | 0.89 | 0.91 |
| | | Alt | 0.89 | 0.91 |
| | | Diff | 0.65 | 0.90 |
| | Objects | Same | 0.88 | 0.91 |
| | | Alt | 0.75 | 0.91 |
| | | Diff | **0.01** | 0.91 |
| | Words | Same | 0.88 | 0.89 |
| | | Alt | 0.88 | 0.89 |
| | | Diff | 0.85 | 0.89 |
| sub-079 | Faces | Same | 0.91 | 0.91 |
| | | Alt | 0.91 | 0.90 |
| | | Diff | 0.91 | 0.92 |
| | Objects | Same | 0.88 | 0.91 |
| | | Alt | 0.91 | 0.91 |
| | | Diff | 0.88 | 0.91 |
| | Words | Same | 0.88 | 0.89 |

| | | | | |
|---|---|---|---|---|
| | | Alt | 0.88 | 0.89 |
| | | Diff | 0.88 | 0.89 |
| sub-089 | Faces | Same | 0.91 | 0.92 |
| | | Alt | 0.91 | 0.92 |
| | | Diff | 0.91 | 0.92 |
| | Objects | Same | 0.88 | 0.91 |
| | | Alt | 0.88 | 0.91 |
| | | Diff | 0.88 | 0.91 |
| | Words | Same | 0.85 | 0.89 |
| | | Alt | 0.88 | 0.89 |
| | | Diff | 0.88 | 0.89 |
| sub-091 | Faces | Same | 0.91 | 0.91 |
| | | Alt | 0.91 | 0.91 |
| | | Diff | 0.91 | 0.91 |
| | Objects | Same | 0.88 | 0.91 |
| | | Alt | 0.88 | 0.91 |
| | | Diff | 0.88 | 0.91 |
| | Words | Same | 0.88 | 0.89 |
| | | Alt | 0.85 | 0.89 |
| | | Diff | 0.85 | 0.89 |
| Preserved RH OTC Patients | | | | |
| sub-066 | Faces | Same | 0.87 | 0.90 |
| | | Alt | 0.87 | 0.90 |
| | | Diff | 0.87 | 0.90 |
| | Objects | Same | 0.88 | 0.91 |
| | | Alt | 0.88 | 0.91 |
| | | Diff | 0.9 | 0.91 |
| | Words | Same | 0.81 | 0.89 |
| | | Alt | 0.88 | 0.89 |
| | | Diff | 0.85 | 0.89 |
| sub-069 | Faces | Same | - | - |
| | | Alt | - | - |
| | | Diff | - | - |
| | Objects | Same | 0.91 | 0.91 |
| | | Alt | 0.91 | 0.91 |
| | | Diff | 0.90 | 0.91 |
| | Words | Same | 0.85 | 0.89 |
| | | Alt | 0.85 | 0.89 |
| | | Diff | 0.88 | 0.89 |

| | | | | |
|---|---|---|---|---|
| sub-090 | Faces | Same | 0.87 | 0.90 |
| | | Alt | 0.91 | 0.90 |
| | | Diff | 0.91 | 0.90 |
| | Objects | Same | 0.88 | 0.91 |
| | | Alt | 0.88 | 0.91 |
| | | Diff | 0.90 | 0.91 |
| | Words | Same | 0.85 | 0.89 |
| | | Alt | 0.85 | 0.89 |
| | | Diff | 0.85 | 0.89 |
| sub-092 | Faces | Same | 0.87 | 0.90 |
| | | Alt | 0.65 | 0.90 |
| | | Diff | 0.65 | 0.90 |
| | Objects | Same | 0.65 | 0.91 |
| | | Alt | 0.88 | 0.91 |
| | | Diff | 0.88 | 0.91 |
| | Words | Same | 0.88 | 0.89 |
| | | Alt | 0.88 | 0.89 |
| | | Diff | 0.91 | 0.89 |
| Preserved LH Control Patients | | | | |
| sub-072 | Faces | Same | 0.91 | 0.92 |
| | | Alt | 0.91 | 0.92 |
| | | Diff | 0.89 | 0.90 |
| | Objects | Same | 0.88 | 0.91 |
| | | Alt | 0.88 | 0.91 |
| | | Diff | 0.88 | 0.91 |
| | Words | Same | 0.85 | 0.89 |
| | | Alt | 0.85 | 0.89 |
| | | Diff | 0.81 | 0.89 |
| sub-073 | Faces | Same | 0.84 | 0.90 |
| | | Alt | 0.81 | 0.90 |
| | | Diff | 0.65 | 0.90 |
| | Objects | Same | 0.91 | 0.91 |
| | | Alt | 0.90 | 0.91 |
| | | Diff | 0.88 | 0.91 |
| | Words | Same | 0.88 | 0.89 |
| | | Alt | 0.88 | 0.89 |
| | | Diff | 0.85 | 0.89 |
| sub-075 | Faces | Same | 0.91 | 0.90 |
| | | Alt | 0.91 | 0.90 |

| | | | | |
|---|---|---|---|---|
| | | Diff | 0.91 | 0.90 |
| | Objects | Same | 0.88 | 0.91 |
| | | Alt | 0.88 | 0.91 |
| | | Diff | 0.88 | 0.91 |
| | Words | Same | - | - |
| | | Alt | - | - |
| | | Diff | - | - |
| sub-080 | Faces | Same | 0.91 | 0.92 |
| | | Alt | 0.91 | 0.91 |
| | | Diff | 0.87 | 0.90 |
| | Objects | Same | 0.88 | 0.91 |
| | | Alt | 0.88 | 0.91 |
| | | Diff | 0.88 | 0.91 |
| | Words | Same | - | - |
| | | Alt | - | - |
| | | Diff | - | - |
| sub-086 | Faces | Same | 0.87 | 0.90 |
| | | Alt | 0.91 | 0.90 |
| | | Diff | 0.91 | 0.90 |
| | Objects | Same | 0.88 | 0.91 |
| | | Alt | 0.88 | 0.91 |
| | | Diff | 0.88 | 0.91 |
| | Words | Same | 0.88 | 0.89 |
| | | Alt | 0.85 | 0.89 |
| | | Diff | 0.81 | 0.89 |
| Preserved RH Control Patients | | | | |
| sub-007 | Faces | Same | 0.91 | 0.90 |
| | | Alt | 0.91 | 0.90 |
| | | Diff | 0.91 | 0.91 |
| | Objects | Same | 0.88 | 0.91 |
| | | Alt | 0.91 | 0.91 |
| | | Diff | 0.88 | 0.91 |
| | Words | Same | - | - |
| | | Alt | - | - |
| | | Diff | - | - |
| sub-045 | Faces | Same | 0.91 | 0.90 |
| | | Alt | 0.91 | 0.90 |
| | | Diff | 0.91 | 0.91 |
| | Objects | Same | 0.90 | 0.91 |

| | | | | |
|---|---|---|---|---|
| | | Alt | 0.90 | 0.91 |
| | | Diff | 0.90 | 0.91 |
| | Words | Same | - | - |
| | | Alt | - | - |
| | | Diff | - | - |
| sub-070 | Faces | Same | 0.91 | 0.90 |
| | | Alt | 0.91 | 0.90 |
| | | Diff | 0.91 | 0.92 |
| | Objects | Same | 0.88 | 0.91 |
| | | Alt | 0.88 | 0.91 |
| | | Diff | 0.91 | 0.91 |
| | Words | Same | 0.88 | 0.89 |
| | | Alt | 0.85 | 0.89 |
| | | Diff | 0.88 | 0.89 |
| sub-076 | Faces | Same | 0.89 | 0.90 |
| | | Alt | 0.89 | 0.90 |
| | | Diff | 0.89 | 0.90 |
| | Objects | Same | - | - |
| | | Alt | - | - |
| | | Diff | - | - |
| | Words | Same | - | - |
| | | Alt | - | - |
| | | Diff | - | - |
| sub-078 | Faces | Same | 0.91 | 0.90 |
| | | Alt | 0.65 | 0.90 |
| | | Diff | 0.06 | 0.50 |
| | Objects | Same | 0.88 | 0.91 |
| | | Alt | 0.91 | 0.91 |
| | | Diff | 0.88 | 0.91 |
| | Words | Same | 0.88 | 0.89 |
| | | Alt | 0.88 | 0.89 |
| | | Diff | 0.85 | 0.89 |
| sub-081 | Faces | Same | 0.91 | 0.90 |
| | | Alt | 0.91 | 0.90 |
| | | Diff | 0.91 | 0.91 |
| | Objects | Same | 0.88 | 0.91 |
| | | Alt | 0.88 | 0.91 |
| | | Diff | 0.89 | 0.91 |
| | Words | Same | 0.85 | 0.89 |

| | | | | |
|---|---|---|---|---|
| | | Alt | 0.85 | 0.89 |
| | | Diff | 0.81 | 0.89 |
| sub-082 | Faces | Same | 0.91 | 0.91 |
| | | Alt | 0.91 | 0.91 |
| | | Diff | 0.91 | 0.92 |
| | Objects | Same | 0.91 | 0.91 |
| | | Alt | 0.9 | 0.91 |
| | | Diff | 0.88 | 0.91 |
| | Words | Same | 0.85 | 0.89 |
| | | Alt | 0.85 | 0.89 |
| | | Diff | 0.85 | 0.89 |

Significant *p*-value is bolded. TD = typically developing controls. LH = left hemisphere. RH = right hemisphere. CS = category selectivity amplitude. RS = repetition suppression magnitude.

**Table S8**: *Model Selection (Repetition Suppression)*

Models

| # | Model | AIC |
|---|---|---|
| 1 | RS ~ group * hemi * stim + age + (1\|ID) | -70.69 |
| 2 | **RS ~ group * hemi + group * stim + hemi * stim + age + (1\|ID)** | -75.06 |
| 3 | RS ~ group * hemi + group * stim + age + (1\|ID) | -70.27 |
| 4 | RS ~ group * hemi + hemi * stim + age + (1\|ID) | -74.82 |
| 5 | RS ~ group * stim + hemi * stim + age + (1\|ID) | -76.61 |

LRTs

| # vs. # | $\chi^2$ | Df | *p* |
|---|---|---|---|
| 1 vs. 2 | 3.63 | 4 | 0.46 |
| 2 vs. 3 | 8.79 | 2 | **0.01** |
| 2 vs. 4 | 8.24 | 4 | 0.08 |
| 2 vs. 5 | 2.45 | 2 | 0.29 |

Selected model and significant *p*-value are bolded. LRT = likelihood ratio test. AIC = Akaike information criterion. Df = degrees of freedom. RS = repetition suppression magnitude. hemi = hemisphere. stim = stimulus category.

**Table S9**: *Post Hoc Comparisons (RS Magnitude ~ Hemisphere x Stimulus Category)*

| Comparison | *z* | *p* |
|---|---|---|
| LH Faces - RH Faces | -0.03 | 0.98 |
| LH Faces - LH Objects | -0.76 | 0.67 |
| LH Faces - LH Words | -3.84 | **< 0.01** |
| RH Faces - RH Objects | -1.29 | 0.44 |
| RH Faces - RH Words | -0.74 | 0.69 |
| LH Objects - RH Objects | -0.50 | 0.73 |
| LH Objects - LH Words | -3.15 | **0.01** |
| RH Objects - RH Words | 0.45 | 0.73 |
| LH Words - RH Words | 2.51 | **0.04** |

Significant *p*-values are bolded. RS = repetition suppression. LH = left hemisphere. RH = right hemisphere.

**Table S10**: *Model Selection (RSA Model Fitting)*

Models

| # | Model | AIC |
|---|---|---|
| 1 | model fit ~ group * hemi * stim + age + (1\|ID) | -96.49 |

| | | |
|---|---|---|
| 2 | model fit ~ group * hemi + group * stim + hemi * stim + age + (1\|ID) | -98.87 |
| 3 | model fit ~ group * hemi + group * stim + age + (1\|ID) | -100.99 |
| 4 | model fit ~ group * hemi + hemi * stim + age + (1\|ID) | -106.76 |
| 5 | model fit ~ group * stim + hemi * stim + age + (1\|ID) | -102.02 |
| 6 | model fit ~ hemi * stim + group + age + (1\|ID) | -109.91 |
| 7 | model fit ~ group * hemi + stim + age + (1\|ID) | -108.91 |
| 8 | model fit ~ group * stim + hemi + age + (1\|ID) | -104.14 |
| 9 | **model fit ~ group + stim + hemi + age + (1\|ID)** | -112.06 |
| 10 | model fit ~ group + hemi + age + (1\|ID) | 166.69 |
| 11 | model fit ~ group + stim + age + (1\|ID) | -110.41 |

LRTs

| # vs. # | $\chi^2$ | Df | *p* |
|---|---|---|---|
| 1 vs. 2 | 5.63 | 4 | 0.23 |
| 2 vs. 3 | 1.88 | 2 | 0.39 |
| 2 vs. 4 | 0.11 | 4 | ~ 1 |
| 2 vs. 5 | 0.85 | 2 | 0.65 |
| 3 vs. 4 | < 0.001 | 2 | ~ 1 |
| 4 vs. 5 | < 0.001 | 2 | ~ 1 |
| 3 vs. 7 | 0.08 | 4 | ~ 1 |
| 3 vs. 8 | 0.85 | 2 | 0.65 |
| 4 vs. 6 | 0.85 | 2 | 0.65 |
| 4 vs. 7 | 1.85 | 2 | 0.40 |
| 5 vs. 6 | 0.11 | 4 | ~ 1 |
| 5 vs. 8 | 1.88 | 2 | 0.39 |
| 6 vs. 8 | < 0.001 | 2 | ~ 1 |
| 7 vs. 8 | < 0.001 | 2 | ~ 1 |
| 6 vs. 9 | 1.85 | 2 | 0.40 |
| 7 vs. 9 | 0.85 | 2 | 0.65 |
| 8 vs. 9 | 0.08 | 4 | ~ 1 |
| 9 vs. 10 | 282.75 | 2 | **< 0.001** |
| 9 vs. 11 | 3.65 | 1 | 0.06 |

Selected model and significant *p*-value are bolded. RSA = representational similarity analysis. LRT = likelihood ratio test. AIC = Akaike information criterion. Df = degrees of freedom. hemi = hemisphere. stim = stimulus category.

**Table S11**: *Crawford Statistics (RSA Model Fitting)*

| ID | Model | *p* (comparison to TD LH) | *p* (comparison to TD RH) |
|---|---|---|---|
| Preserved LH OTC Patients | | | |
| sub-004 | CS | 0.86 | 0.68 |
| | RS | 0.85 | 0.74 |
| | CS-RS | 0.87 | 0.71 |
| sub-077 | CS | 0.86 | 0.68 |
| | RS | 0.85 | 0.79 |
| | CS-RS | 0.87 | 0.71 |
| sub-079 | CS | 0.86 | 0.78 |
| | RS | 0.85 | 0.74 |
| | CS-RS | 0.87 | 0.83 |
| sub-089 | CS | 0.86 | 0.68 |
| | RS | 0.25 | 0.71 |

| | | | |
|---|---|---|---|
| | CS-RS | 0.87 | 0.71 |
| sub-091 | CS | 0.86 | 0.78 |
| | RS | 0.85 | 0.74 |
| | CS-RS | 0.87 | 0.83 |
| Preserved RH OTC Patients | | | |
| sub-066 | CS | 0.86 | 0.68 |
| | RS | 0.2 | 0.23 |
| | CS-RS | 0.87 | 0.71 |
| sub-069 | CS | 0.86 | 0.78 |
| | RS | 0.85 | 0.74 |
| | CS-RS | 0.87 | 0.83 |
| sub-090 | CS | 0.86 | 0.85 |
| | RS | 0.61 | 0.74 |
| | CS-RS | 0.87 | 0.85 |
| sub-092 | CS | 0.86 | 0.85 |
| | RS | 0.2 | 0.39 |
| | CS-RS | 0.87 | 0.85 |
| Preserved LH Control Patients | | | |
| sub-072 | CS | 0.86 | 0.85 |
| | RS | 0.85 | 0.79 |
| | CS-RS | 0.87 | 0.83 |
| sub-073 | CS | 0.86 | 0.68 |
| | RS | 0.85 | 0.74 |
| | CS-RS | 0.87 | 0.71 |
| sub-075 | CS | 0.86 | 0.68 |
| | RS | 0.31 | 0.74 |
| | CS-RS | 0.87 | 0.71 |
| sub-080 | CS | 0.86 | 0.68 |
| | RS | **0.01** | 0.12 |
| | CS-RS | 0.87 | 0.71 |
| sub-086 | CS | 0.86 | 0.68 |
| | RS | 0.85 | 0.74 |
| | CS-RS | 0.87 | 0.71 |
| Preserved RH Control Patients | | | |
| sub-007 | CS | 0.86 | 0.68 |
| | RS | 0.85 | 0.82 |
| | CS-RS | 0.87 | 0.71 |
| sub-045 | CS | 0.86 | 0.78 |

| | | | |
|---|---|---|---|
| | RS | 0.85 | 0.74 |
| | CS-RS | 0.87 | 0.83 |
| sub-070 | CS | 0.86 | 0.68 |
| | RS | 0.20 | 0.62 |
| | CS-RS | 0.87 | 0.71 |
| sub-076 | CS | 0.86 | 0.68 |
| | RS | 0.21 | 0.23 |
| | CS-RS | 0.87 | 0.71 |
| sub-078 | CS | 0.86 | 0.78 |
| | RS | 0.61 | 0.71 |
| | CS-RS | 0.87 | 0.83 |
| sub-081 | CS | 0.86 | 0.78 |
| | RS | 0.85 | 0.74 |
| | CS-RS | 0.87 | 0.83 |
| sub-082 | CS | 0.86 | 0.78 |
| | RS | 0.85 | 0.74 |
| | CS-RS | 0.87 | 0.83 |

Significant *p*-value is bolded. TD = typically developing controls. LH = left hemisphere. RH = right hemisphere. CS = category selectivity. RS = repetition suppression.

**Table S12**: *Model Selection (Behavior)*

| Models | | |
|---|---|---|
| # | Model | AIC |
| 1 | **accuracy ~ group * stim + age + (1\|ID)** | -181.60 |
| 2 | accuracy ~ group + stim + age + (1\|ID) | -174.36 |

| LRTs | | | |
|---|---|---|---|
| # vs. # | $\chi^2$ | Df | *p* |
| 1 vs. 2 | 15.25 | 4 | **< 0.01** |

Selected model and significant *p*-value are bolded. LRT = likelihood ratio test. AIC = Akaike information criterion. Df = degrees of freedom. stim = stimulus category.

**Table S13**: *Post Hoc Comparisons (Behavioral Accuracy ~ Group x Stimulus Category)*

| Stimulus Category | Comparison | *z* | *p* | *p* (permutation testing) |
|---|---|---|---|---|
| Faces | OTC - TD | -4.06 | **< 0.001** | **< 0.001** |
| | OTC - CP | -3.67 | **< 0.01** | **< 0.001** |
| | TD - CP | 0.10 | 0.92 | 0.93 |
| Objects | OTC - TD | -5.37 | **< 0.001** | **< 0.001** |
| | OTC - CP | -3.45 | **< 0.01** | **0.03** |
| | TD - CP | 2.02 | 0.08 | 0.27 |
| Words | OTC - TD | -0.90 | 0.55 | 0.27 |
| | OTC - CP | -0.70 | 0.62 | 0.31 |
| | TD - CP | 0.14 | 0.92 | 0.93 |

Significant *p*-values are bolded. OTC = occipitotemporal cortex patients. TD = typically developing controls. CP = control patients.

**Table S14**: *Model Selection (Behavior - Patient Data Only)*

| Models | | |
|---|---|---|
| # | Model | AIC |
| 1 | accuracy ~ group * hemi * stim + age + (1\|ID) | -79.57 |
| 2 | **accuracy ~ group * hemi + group * stim + hemi * stim + age + (1\|ID)** | -80.79 |
| 3 | accuracy ~ group * hemi + group * stim + age + (1\|ID) | -84.58 |
| 4 | accuracy ~ group * hemi + hemi * stim + age + (1\|ID) | -78.74 |
| 5 | accuracy ~ group * stim + hemi * stim + age + (1\|ID) | -78.77 |

| LRTs | | | |
|---|---|---|---|
| # vs. # | $\chi^2$ | Df | *p* |
| 1 vs. 2 | 2.78 | 2 | 0.25 |
| 2 vs. 3 | 4.21 | 4 | 0.38 |
| 2 vs. 4 | 6.05 | 2 | **0.05** |
| 2 vs. 5 | 4.03 | 1 | **0.05** |

Selected model and significant *p*-value are bolded. LRT = likelihood ratio test. AIC = Akaike information criterion. Df = degrees of freedom. hemi = hemisphere. stim = stimulus category.

**Table S15**: *Post Hoc Comparisons (Behavioral Accuracy ~ Group x Hemisphere) (Patient Data Only)*

| Comparison | *z* | *p* |
|---|---|---|
| CP LH - OTC LH | 0.80 | 0.42 |
| CP LH - OTC RH | -1.58 | 0.20 |
| OTC LH - OTC RH | 1.44 | 0.20 |
| CP RH - OTC RH | 3.83 | **< 0.01** |

Significant *p*-value is bolded. OTC = occipitotemporal cortex patients. CP = control patients. LH = left hemisphere. RH = right hemisphere.

**Table S16**: *Karolinska Directed Emotional Faces[1] Stimulus IDs*

| |
|---|
| AM02NES |
| AM03NES |
| AM05NES |
| AM06NES |
| AM07NES |
| AM10NES |
| AM11NES |
| AM13NES |
| AM17NES |
| AM18NES |
| AM21NES |
| AM23NES |
| AM25NES |
| AM26NES |
| AM27NES |
| AM31NES |
| AM32NES |
| AM35NES |

**Table S17**: *R Packages*

| |
|---|
| broom[2] |
| car[3] |

| |
|---|
| lme4[4] |
| nnet[5] |
| pracma[6] |
| psych[7] |
| singcar[8] |
| strex[9] |
| tidyverse[10] |

### References


1. Lundqvist, D., Flykt, A. & Öhman, A. *The Karolinska Directed Emotional Faces - KDEF*. (CD ROM from Department of Clinical Neuroscience, Psychology section, Karolinska Institutet, ISBN 91-630-7164-9, 1998).
2. Robinson, D. broom: An R Package for Converting Statistical Analysis Objects Into Tidy Data Frames. *arXiv* (2014).
3. Companion to Applied Regression [R package car version 3.0-12]. (2021). at <https://cran.r-project.org/web/packages/car/index.html>
4. Bates, D., Mächler, M., Bolker, B. & Walker, S. Fitting linear mixed-effects models using lme4. *J. Stat. Softw.* **67,** 1–48 (2015).
5. Venables, W. N. & Ripley, B. D. *Modern Applied Statistics with S (Statistics and Computing)*. 510 (Springer, 2002).
6. Borchers, H. W. *pracma: Practical Numerical Math Functions*. (2021).
7. Revelle, W. *psych: Procedures for Personality and Psychological Research*. (Northwestern Unviersity, 2021).
8. Rittmo, J. & McIntosh, R. *singcar: Comparing Single Cases to Small Samples*. (2021).
9. Nolan, R. *strex: Extra String Manipulation Functions*. (2021).
10. Wickham, H. *et al.* Welcome to the tidyverse. *JOSS* **4,** 1686 (2019).
